# Neuronal apoptosis drives remodeling states of microglia and shifts in survival pathway dependence

**DOI:** 10.1101/2022.01.05.475126

**Authors:** Sarah R Anderson, Jacqueline M Roberts, Nate Ghena, Emmalyn Irvin, Joon Schwakopf, Isabelle Cooperstein, Alejandra Bosco, Monica L Vetter

## Abstract

Microglia serve critical remodeling roles that shape the developing nervous system, responding to the changing neural environment with phagocytosis or soluble factor secretion. Recent single-cell sequencing (scRNAseq) studies have revealed the context-dependent diversity in microglial properties and gene expression, but the cues promoting this diversity are not well defined. Here, we ask how interactions with apoptotic neurons shape microglial state, including lysosomal and lipid metabolism gene expression and independence from Colony-stimulating factor 1 receptor (CSF1R) for survival. Using early postnatal mouse retina, a CNS region undergoing significant developmental remodeling, we performed scRNAseq on microglia from mice that are wild-type, lack neuronal apoptosis (Bax KO), or are treated with CSF1R inhibitor (PLX3397). We find that interactions with apoptotic neurons drives multiple microglial remodeling states, subsets of which are resistant to CSF1R inhibition. We find that TAM receptor Mer and complement receptor 3 are required for clearance of apoptotic neurons, but that Mer does not drive expression of remodeling genes. We show TAM receptor Axl is negligible for phagocytosis or remodeling gene expression but is consequential for microglial survival in the absence of CSF1R signaling. Thus, interactions with apoptotic neurons shift microglia towards distinct remodeling states and through Axl, alters microglial dependence on survival pathway, CSF1R.

## Introduction

Microglia are parenchymal innate immune cells and vital remodelers of the central nervous system (CNS) [1]. They have a myriad of critical functions during development, including elimination of viable or dying cells [2]. Similar to other macrophages, microglia are extremely attuned to changes in their neural niche [3–5] and rapidly respond by phagocytosis or secreting soluble factors. Recent single-cell RNA sequencing (scRNAseq) has demonstrated that microglia are particularly diverse during development [6, 7]. However, we still lack an understanding of the drivers of this heterogeneity, including the impact of environmental cues or the act of phagocytosis itself. Therefore, connecting microglial states to specific developmental events and determining the pathways involved remains a central challenge.

One developmental process fundamental for nearly every organ in the body is apoptotic death of excess or dysfunctional cells [8]. In the CNS, death of neurons and glia is an essential and well-documented feature of development [9]. Efferocytosis, or the phagocytosis of apoptotic cells, is a rapid and carefully orchestrated process designed to minimize damage to surrounding cells [10, 11]. Largely mediated by microglia, efferocytosis is important for maintaining CNS homeostasis not only in development but in aging and disease as well [12, 13]. The developing mouse retina is an excellent CNS model system to link remodeling events such as efferocytosis and microglial state and function [14]. During the postnatal period in the retina, as circuits are being established, waves of neuronal death occur [15]. For example, retinal ganglion cells (RGCs), the projection neurons connecting retina and brain, undergo a well-characterized culling during the first postnatal week which is dependent on pro-apoptotic factor, Bcl-2- associated X protein (Bax) [16]. An estimated ∼50% of RGCs will undergo apoptosis and need to be cleared [17, 18]. We previously found that neuronal death significantly influences microglia properties in postnatal retina, resulting in a majority of microglia expressing a distinct gene signature and showing independence from colony-stimulating factor 1 receptor (CSF1R) signaling for survival [19]. Relative to adult microglia, homeostatic genes were reduced while genes associated with phagocytosis and lipid metabolism were increased, similar to aging and disease-associated microglia (DAM) [20–22], and to developmental microglia residing in postnatal white matter tracts (PAM) [7] or (ATM) [6] or CD11c microglia [23].

Consistent with either apoptotic cell recognition or phagocytosis playing a central role in driving this gene signature, microglia in disease states increase select DAM genes in response to apoptotic cells [22], and ATM/PAM populations in the white matter tract engulf oligodendrocyte precursor cells [7, 24]. In disease, the TREM2 receptor is partially required for acquisition of the DAM signature [21, 22], but not for the developmental signature in retina [19] or brain [7]. Thus, it remains unclear to what extent the process of phagocytosis is altering properties of developmental microglia or whether distinct recognition pathways are involved. Microglia, like other macrophages, can recognize and engulf apoptotic (or viable) cells via a variety of ligand-receptor systems including C1q/C3 to complement receptor 3 (CR3) and exposed phosphatidylserine (PtdSer) to TAM family of tyrosine kinase receptors Mer (gene Mertk) and Axl [25], but how these pathways mediate changes in microglia properties and responses is an active area of research.

Here, we ask how neuronal apoptosis drives key properties of developmental microglia such as CSF1R independence and expression of DAM/ATM/PAM-related genes, and test the role of phagocytosis and specific recognition pathways. Through scRNAseq on postnatal retinal microglia we find considerable transcriptional heterogeneity, with multiple distinct microglial populations expressing DAM/ATM/PAM-related, lysosomal, and lipid metabolism genes. We further show, by analyzing microglia from Bax knockout (KO) retinas, that exposure to dying neurons drives several of these states and that most are more resistant to CSF1R inhibition. We find that CR3 and Mer are required for clearance of dying RGCs, but not for augmenting microglial survival in the absence of CSF1R signaling. Conversely, Axl is not required for clearance of dying RGCs, but for mediating independence from CSF1R signaling for survival. Loss of Mer did not have a widespread impact on expression of microglial remodeling genes, and neither did loss of Axl or Mertk/Axl double KO (dKO), suggesting that TAM receptor-mediated signaling or reduced clearance are not sufficient. Thus, we find that interactions with apoptotic neurons drives developmental microglial diversity, and that distinct recognition receptors mediate phagocytosis of dying cells versus specific microglial properties, including microglial survival.

## Results

### Multiple microglial states coexist in postnatal retina

To better understand the influence of environmental cues on microglial states, we first sought to understand the extent of microglial heterogeneity in the postnatal retina and then explicitly determine states that were driven by neuronal apoptosis. Second, we previously linked CSF1R independence to a DAM/ATM/PAM-related signature and neuronal apoptosis [19], so we wanted to identify and characterize microglial states that remained following CSF1R inhibition (PLX3397, PLX). Therefore, we performed scRNAseq on sorted retinal microglia from 4 groups at postnatal day 6 and 7 (P6/P7): Bax WT, Bax KO, CX3CR1-GFP/+ given vehicle (daily for 3 days), and CX3CR1-GFP/+ pups dosed with PLX3397 (daily for 3 days) (Figure 1A). Microglia from Bax WT and littermate KOs (CD45^+^ CD11b^+^/CX3CR1-GFP^+^ CCR2^-^) as well as microglia from PLX and vehicle controls (CD45^+^ CX3CR1-GFP^+^ Ly6C^-^) were sorted (Supplemental 1A-D) and sequenced using the 10X Genomics platform. Following sequencing and manual filtering (Supplemental 2), we used in silico Bax genotyping to sort out and reassign a subset of cells from animals incorrectly assigned to the Bax WT and KO groups (Supplemental 3).

**Figure 1.**
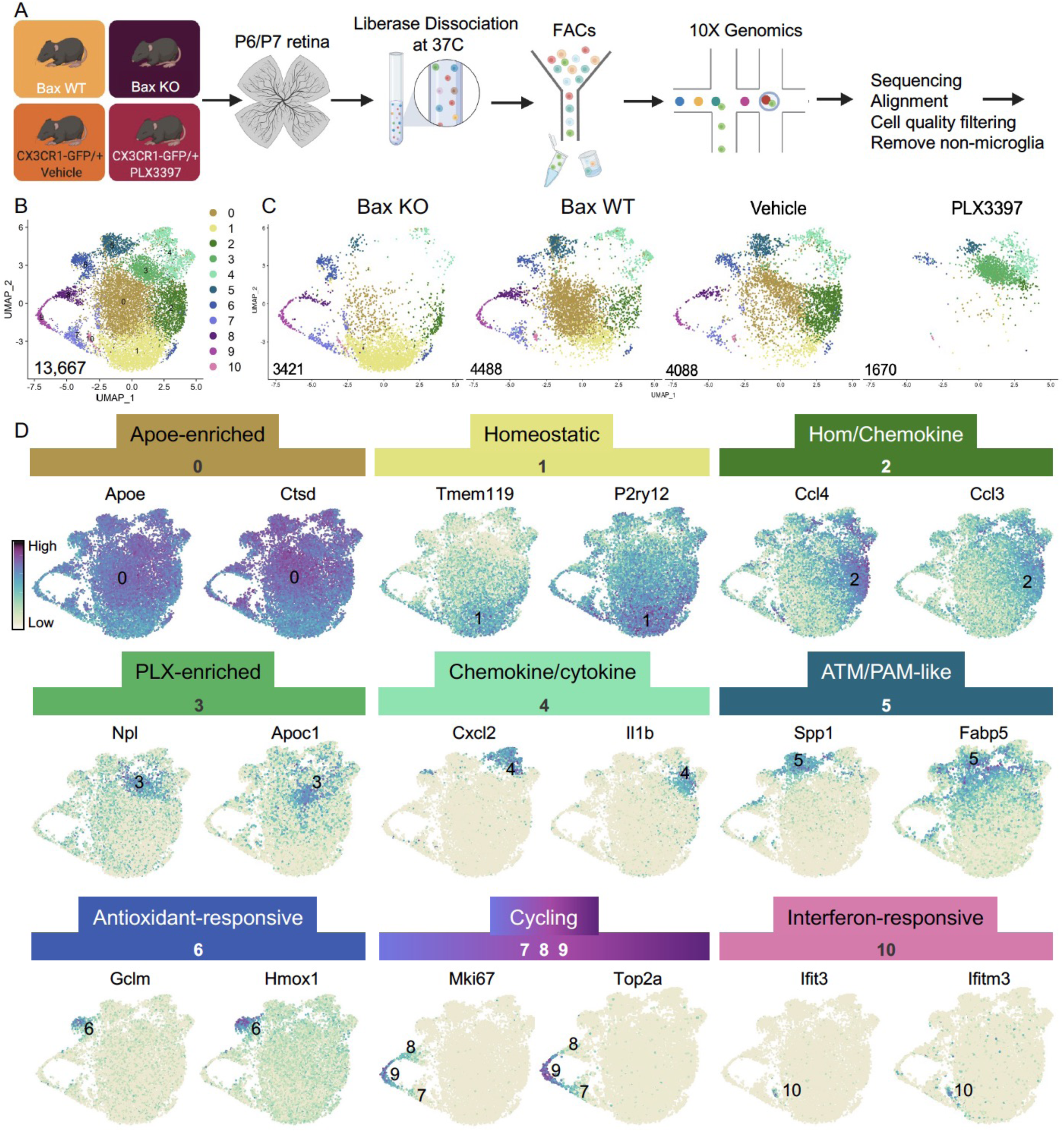
Multiple microglial states coexist in postnatal retina. (A) Workflow for collection, dissociation, sorting, and sequencing of individual microglia from 4 different groups: Bax WT and KO, PLX and vehicle. (B) UMAP plot of 13,667 microglia cells from all 4 samples distributed into 11 clusters by unsupervised clustering. (C) UMAP plots illustrating the distribution of cells for each sample. Number of cells per sample labeled in lower left. (D) UMAP plots of two genes enriched in each cluster. Color scale is based on relative gene expression: dark purple = highest, light yellow = lowest.

Unsupervised clustering was performed on the 15,084 cells from the four groups (4,902 - Bax WT; 4,224 - Bax KO; 4,198 - Vehicle; 1,760 - PLX3397) (Supplemental 4A). Of the 15,084 cells, 1,417 in satellite clusters were deemed non-microglia cells based on expression of established markers and were excluded from further analysis (Supplemental 4B-E). Differences in cell surface markers used to sort Bax WT/KO (CD45^+^CD11b^+^CX3CR1^+^CCR2^-^) compared to PLX/Vehicle (CD45^+^CX3CR1^+^Ly6C^-^) (Supplemental 1) led to variance in the proportion of macrophage/monocyte-like cells (Supplemental 4F), but this had no major impact on microglial cell number (Gray dots, Supplemental 4C,F).

Unsupervised re-clustering of 13,667 remaining cells yielded 11 microglia clusters (Figure 1B). Cells from controls (Bax WT and Vehicle) were represented in every cluster (Figure 1C), indicating that microglia in an individual developing CNS region during a specific time window are strikingly diverse. Further, we found that Bax KO and PLX-treated microglia had dramatic and opposing shifts in their distribution compared to controls (Figure 1C). To determine the characteristics of the 11 clusters, we examined the top genes for each (Figure 1D, Supplemental Table 1). While virtually every cell expressed *Apoe* and *Ctsd*, consistent with our previous findings [19], both genes were significantly enriched in cluster 0, which had high expression of other DAM/PAM/ATM genes such as *Lyz2* and *Cd9*, but also intermediate levels of homeostatic gene expression such as *P2ry12* and *Tmem119*. Thus, we called this cluster Apoe-enriched. Cluster 1 had highest expression of *Tmem119* and *P2ry12* as well as other homeostatic genes, including *Itgam, Siglech, Tgfbr1, P2ry13, Selplg, and Adgrg1* and so we named it Homeostatic. Cluster 2 was enriched for chemokines including *Ccl4* and *Ccl3,* two genes previously found in injury-responsive microglia in demyelinated lesions [6] and in PAM of developing WM tracts [7], and had intermediate/high expression of homeostatic genes (e.g. *Tmem119*; Figure 1D) and so we termed it Hom/chemokine cluster. Since *Ccl3* and *Ccl4* can be induced by dissociation [26] we confirmed expression of *Ccl3* by *in situ* hybridization in whole mount retina (Supplemental 5). Cluster 3 was marked by *Npl*, found in lipid-droplet microglia [27], and *Apoc1*, a lipoprotein high in MS-associated human microglia [28]. Since this cluster was substantially increased in PLX-treated retinas, we named this cluster PLX-enriched. Chemokines *Cxcl2* and *Cxcl10* and pro-inflammatory cytokines, such as *Il1b* and *Tnf,* were high in Cluster 4, dubbed the Chemokine/cytokine cluster. Cluster 5 most resembled ATM [6] (43/68 genes shared) and PAM [7] (41/68 genes shared) with specific expression of *Spp1* and high expression of *Fabp5, Ctsl, Lpl, Igf1,* and *Csf1* and thus we termed it ATM/PAM-like. Cluster 6, was named Antioxidant-responsive cluster due to high expression of antioxidant responsive genes, *Hmox1* and *Gclm* [29]. Clusters 7, 8, and 9 represented Cycling microglia and expressed *Mki67, Top2a*, and *Mcm* genes. Cluster 10, the smallest, contained 71 cells with high and very specific expression of interferon response genes *Ccl5, Ifit3* and *Ifitm3* [30] and was thus termed Interferon-responsive cluster. Thus, these data suggest that within the developing retina, several microglial states coexist including homeostatic and various subsets of DAM/PAM/ATM-related and chemokine-expressing microglia.

### Postnatal retinal microglia encompass a spectrum of homeostatic to remodeling states

We noted a continuum of expression of the homeostatic genes *Tmem119* and *P2ry12* across the clusters that appeared to be inverse to broadly expressed DAM/PAM/ATM genes such as *Apoe, Ctsd* (Figure 1D). We more closely examined the expression of homeostatic genes, as well as genes associated with lysosomal function, lipid metabolism and transport, and other DAM/PAM/ATM-related genes. As for *Tmem119* and *P2ry12*, we found a gradient of expression of other homeostatic genes including *Siglech, Tgfbr1, Mafb* and *Selplg* which were highest in Homeostatic cluster (1), high/intermediate in Hom/chemokine (2), intermediate in Apoe-enriched (0), Chemokine/cytokine cluster (4), ATM/PAM-like (5), and Antioxidant-responsive (6), with lowest expression in PLX-enriched cluster (3), suggesting a spectrum from more to less homeostatic (Figure 2B).

**Figure 2.**
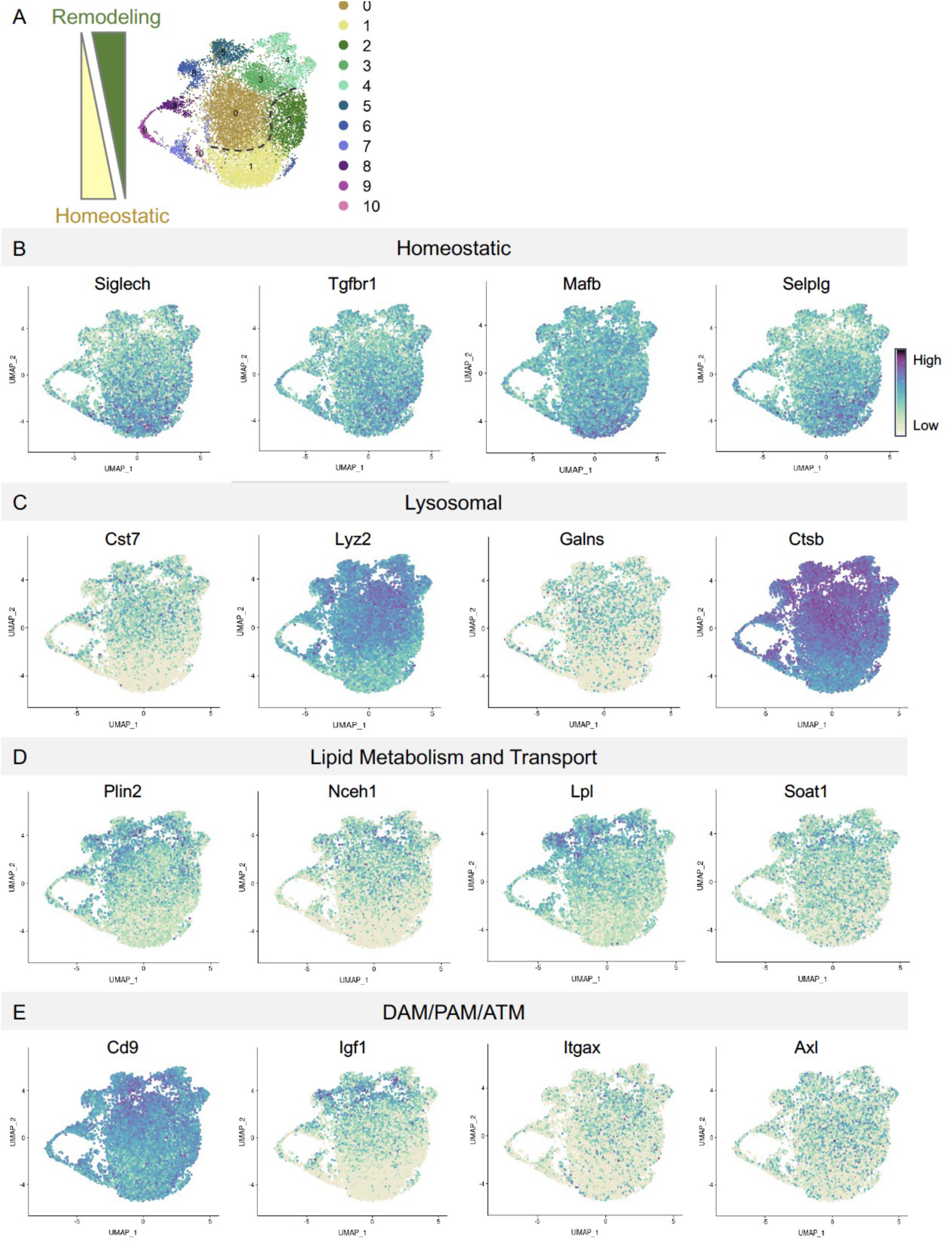
Postnatal retinal microglia encompass a spectrum of homeostatic to remodeling states. (A) UMAP plot of 11 clusters from all sequenced microglia. Spectrum of homeostatic (bottom) to more remodeling (top). Dashed line demarcates in interface between relatively more homeostatic and remodeling populations. (B) UMAP plots of microglial homeostatic genes. (C) UMAP plots of microglial homeostatic genes important for lysosomal function (D) UMAP plots of genes associated with lipid metabolism. (E) UMAP plots of DAM/PAM/ATM genes. Color scale is based on relative gene expression: dark purple = highest, light yellow = lowest.

To determine whether this related to shifting phagocytic function of microglia, we examined the expression of lysosomal genes *Cst7, Lyz2, Galns* and *Ctsb* and saw roughly the opposite pattern. These genes were either absent or sparse in Homeostatic cluster (1) and Hom/chemokine (2) and more highly expressed in Apoe-enriched (0), PLX-enriched (3), Chemokine/cytokine cluster (4), ATM/PAM-like (5), and Antioxidant-responsive (6) (Figure 2C). Notably, *Lyz2* showed highest levels in PLX-enriched (3) the least homeostatic cluster. Additionally, genes involved with lipid metabolism and cholesterol transport, important processes following the engulfment of apoptotic cells [31], followed the same trend (Figure 2D). Thus, microglia in clusters (0,3,4,5 and 6) are equipped to be actively breaking down phagocytosed material compared to more homeostatic clusters (1,2). Therefore, we conclude that clusters 0,3,4,5, and 6 are involved in active remodeling activities and will refer to them as remodeling clusters (Figure 2A). Since we saw enrichment of *Apoe* and *Ctsd* in these remodeling clusters, we examined the expression of additional DAM/ATM/PAM-related genes, including *Cd9, Igf1*, as well as the recognition receptors *Itgax* and *Axl*. We noted similar low expression in Homeostatic cluster (1) and Hom/chemokine (2) and high expression in the remodeling clusters (0,3,4,5,6) (Figure 2E). Interestingly, DAM/PAM/ATM-related genes had varying cluster specificity. Genes such as *Apoe* were expressed in nearly every cell, whereas *Igf1* and *Spp1* were more restricted to specific clusters, suggesting that regulation of these genes is complex (Figure 1D, 2E). Together, these results suggest that phagolysosomal function defines the spectrum of microglial states in the postnatal retina.

### Neuronal apoptosis drives multiple remodeling states

We previously showed that neuronal death has a major influence on expression of remodeling genes in postnatal retinal microglia [19] and wondered whether recognition or clearance of dying neurons was a key factor in driving diverse microglial remodeling states. We examined microglia from Bax KO retinas in which apoptotic death programs in RGCs and other neurons are selectively lost [32] and saw a dramatic shift in cluster distribution with the loss of Bax compared to littermate controls (Figure 3A). We found a 5-fold expansion in the Homeostatic cluster (1) in Bax KO compared to WT (Figure 3A,B,C). This was concurrent with decreases in remodeling clusters Apoe-enriched (0), Chemokine/cytokine expressing (4), and ATM/PAM-like (5), and the minor PLX-enriched cluster (3), which were 4.7-fold, 3.46-fold, 5.86-fold, and 5-fold more abundant in WT, respectively (Figure 3A,B). Clusters that remained largely unaltered included Hom/chemokine (2), Antioxidant-responsive (6), Cycling (7-9), and Interferon-responsive (10), suggesting they were not driven by apoptotic cell interactions or clearance (Figure 3A,B). Therefore, we conclude that the spectrum of homeostatic to more remodeling clusters (0,3,4 and 5) in the postnatal retina is driven predominately by neuronal apoptosis.

**Figure 3.**
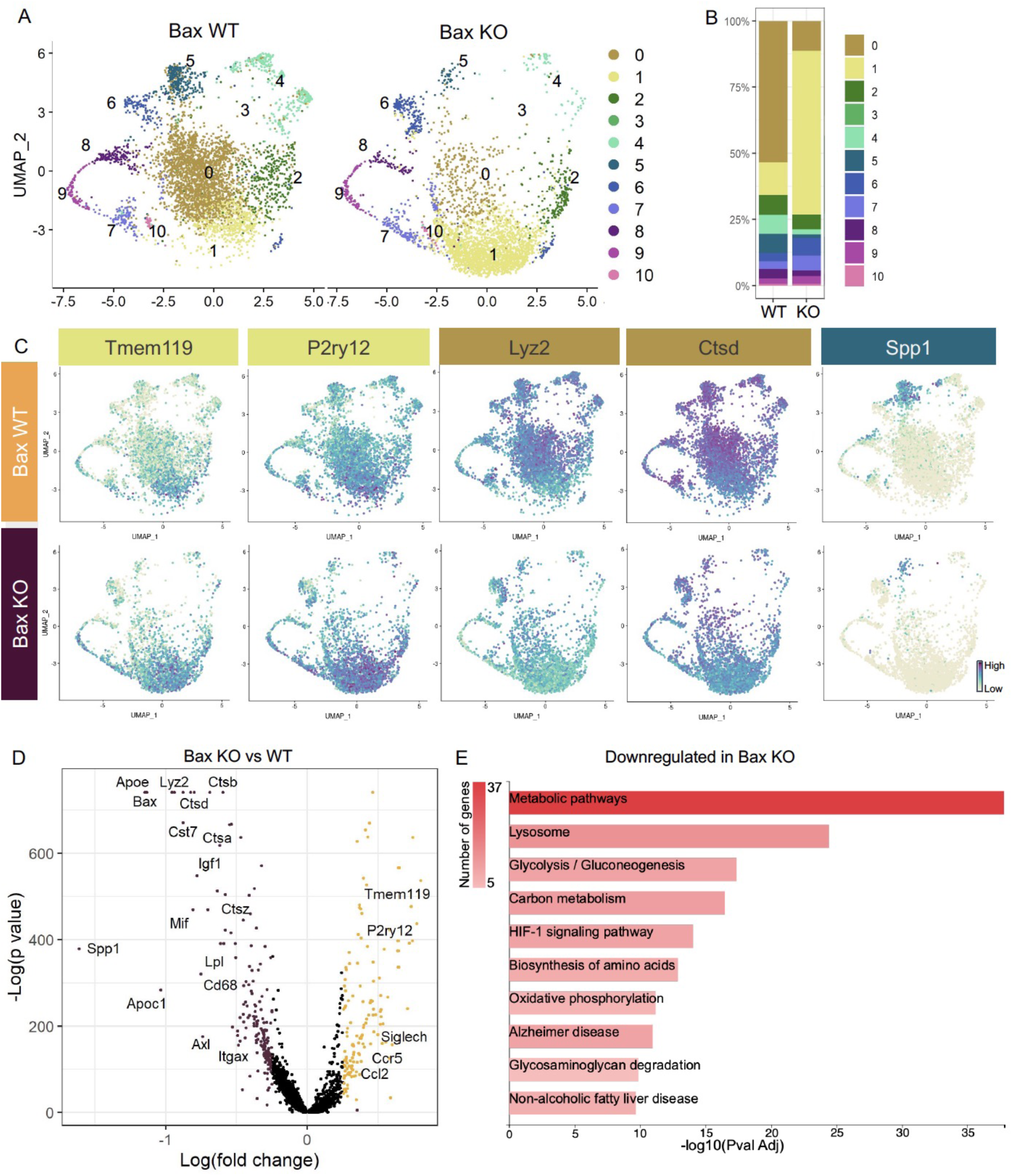
Neuronal death drives multiple microglial remodeling states. (A) UMAP plot of microglia cells from BAX WT (left) and BAX KO (right) samples distributed into 11 clusters. (B) Bar graph of the proportion of cells in each cluster for each sample. (C) UMAP plots showing expression of representative genes from selected clusters. (D) Volcano plot showing differential gene expression of all BAX KO cells compared to BAX WT cells. Each gene is plotted according to the significance (-Log(p value)) and magnitude (Log(fold change)) of the difference such that those genes enriched in BAX KO have Log(fold change) > 0. Red points indicate genes with p-value ≤0.05 and Log(fold change) > 0.25. (E) KEGG pathway analysis of 183 genes downregulated in BAX KO compared to WT using GeneCodis 4.

To further examine microglial genes dependent on neuronal apoptosis, we compared all cells from each genotype, ignoring cluster membership, and identified genes that were specific to each sample (Figure 3D, Supplemental Table 2). Consistent with a shift to a more homeostatic state, *Tmem119, P2ry12*, and *Siglech* were among the 133 upregulated genes in microglia from Bax KO retina. The 183 downregulated genes included lysosomal and lipid metabolism genes such as *Apoe, Lyz2, Cst7, Ctsb, Lpl*, and *Cd68,* DAM/PAM/ATM-related genes such as *Igf1* and *Spp1,* and recognition receptors *Itgax* and *Axl* (Figure 3C,D). We next performed KEGG analysis using GeneCodis 4 [33] on downregulated genes in Bax KO to identify major pathways driven by interactions with dying neurons (Figure 3E). Consistent with the idea that microglial phagocytosis was a key factor in these gene expression changes, we found metabolic and lysosomal pathways significantly reduced. To determine whether loss of Bax altered microglia density or distribution, we used C1q as a marker for microglia, which we validated by both scRNAseq (Supplemental 4C) and immunostaining (Supplemental 6A,B). We found that microglia remained uniformly spaced but that density was reduced by nearly half in Bax KO retinas (Supplemental 6C,D), consistent with our prior flow cytometry analysis [19]. Altogether, this suggests that neuronal apoptosis regulates important properties of microglia in the postnatal retina, including overall density as well as the emergence of multiple remodeling states.

### Subsets of remodeling states survive CSF1R inhibition, while homeostatic microglia are more vulnerable

A striking property of microglia in postnatal retina is that ∼60% survive inhibition of CSF1R signaling [19], while only very small subsets show this property in adult brain [34]. We previously found that surviving microglia had reduced homeostatic gene expression and increased DAM/PAM/ATM gene expression, and this required neuronal apoptosis [19]. Since we found that the less homeostatic remodeling clusters (0,3,4,5) were driven by neuronal apoptosis, we investigated whether these microglial states would be more resistant to CSF1R inhibition. We examined scRNAseq of Vehicle and PLX-treated microglia (dosed for three days) and found a large shift in the distribution of cells across clusters with PLX treatment (Figure 4A). Several remodeling clusters were either enriched or maintained. The least homeostatic cluster, PLX- enriched (3), which was a very small proportion of cells in both Vehicle and Bax WT (Figure 3A,B), represented nearly 60% of remaining microglia following PLX3397 (increased 60-fold) and thus seemed to arise, in part, by treatment (Figure 4A,B). This could potentially be due to the further reduction of homeostatic properties of surviving microglia with CSF1R blockade [35]. Remodeling cluster Chemokine/cytokine (4) was increased 2-fold with PLX-treatment and ATM/PAM-like (5), Antioxidant-responsive (6), and Interferon-responsive (10) were slightly reduced or unchanged (0.755-fold, 0.654-fold, 0.979-fold, respectively), suggesting they are more resistant than other subsets (Figure 4A,B). Conversely, we found clusters that had high or intermediate expression of homeostatic genes, Apoe-enriched (0), Homeostatic (1), Hom/chemokine (2), and Cycling (7,8,9) were largely absent following PLX treatment, suggesting these clusters are more dependent on CSF1R for survival or that they had shifted to a different state (Figure 4A,B). We noted the overlap between clusters dependent on neuronal apoptosis (0,3,4 and 5) and resistant to loss of CSF1R signaling (3,4,5,6 and 10), illustrating the link between the two and further arguing that certain microglial states driven by neuronal apoptosis confer CSF1R independence. Furthermore, we found that downregulation of *Csf1r* alone does confer independence as more resistant clusters (3,4,5 and 6) had slightly reduced expression of *Csf1r* compared to susceptible clusters (0,1, and 2) but comparable to other susceptible clusters (7,8,9) (Figure 4D), consistent with our previous findings [19].

**Figure 4.**
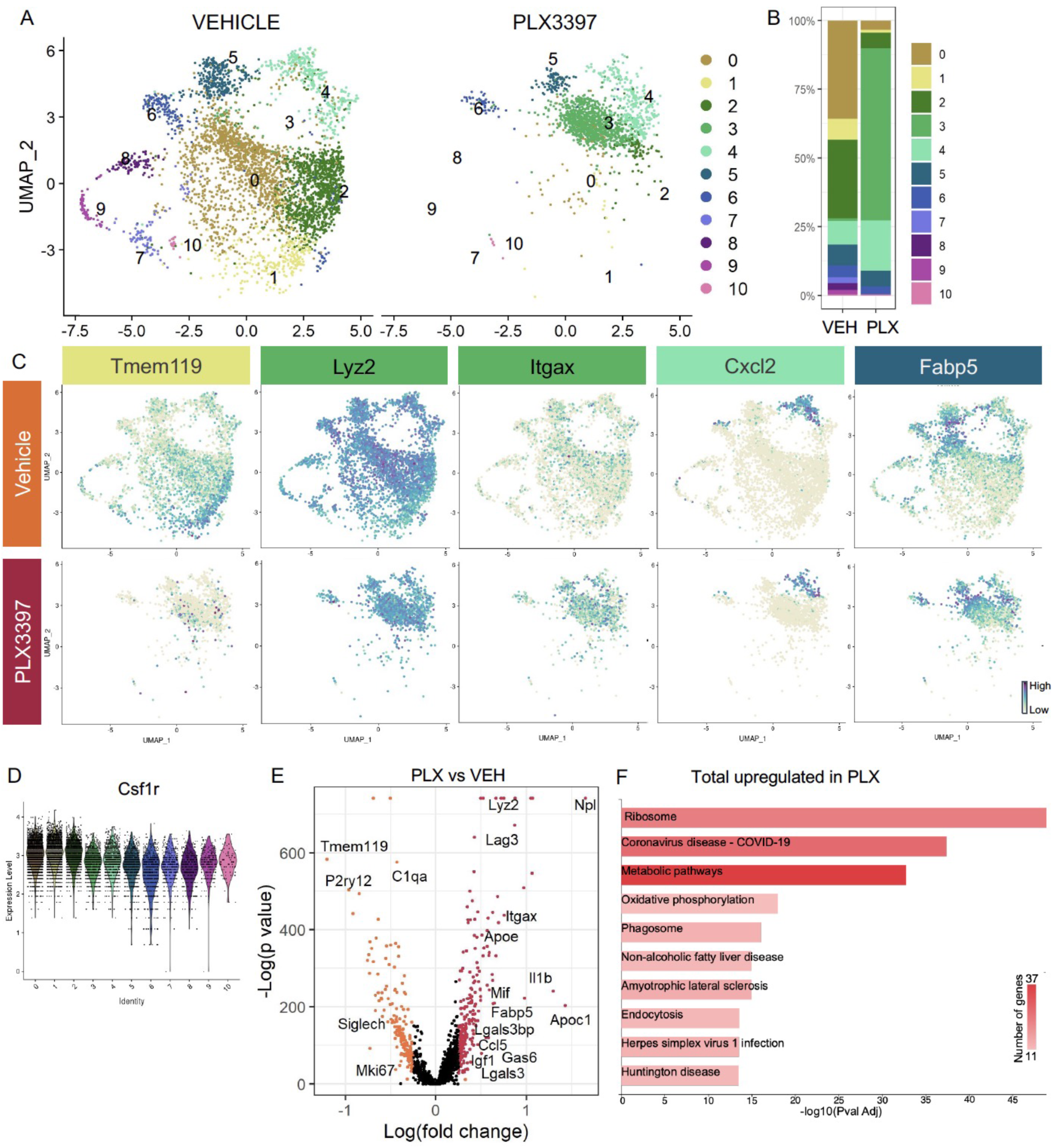
Subsets of remodeling states survive CSF1R inhibition, while homeostatic microglia are more vulnerable. (A) UMAP plot of microglia cells from Vehicle (left) and PLX3397 (right) samples distributed into 11 clusters. (B) Proportion of cells from each sample within each cluster. (C) UMAP plots showing expression of representative genes from selected clusters. (D) Violin plot of Csf1r expression across clusters. (E) Volcano plot showing differential gene expression of pseudobulked PLX cells compared to Vehicle cells. Each gene is plotted according to the significance (-Log(p value)) and magnitude (Log(fold change)) of the difference such that those genes enriched in PLX have Log(fold change) > 0. Red points indicate genes with p-value ≤ 0.05 and Log(fold change) > 0.25. (F) KEGG Pathway analysis of 254 upregulated genes in PLX compared to Vehicle using GeneCodis 4.

To further probe genes and pathways enriched following CSF1R inhibition, we performed a pairwise comparison on all Vehicle and PLX-treated cells, regardless of cluster. We found that DAM/ATM/PAM-related genes *Apoe, Itgax,* and *Fabp5*, lysosomal genes *Lyz2* and *Ctss,* and lipid metabolism genes *Nceh1*, *Soat1, Abca1,* and *Apoc1* were among the 254 upregulated genes in PLX-treated cells (Figure 4C,E, Supplemental Table 3). Consistent with a loss of more homeostatic microglia, homeostatic genes including *Tmem119, P2ry12,* and *Siglech* were significantly reduced (181 genes downregulated) (Figure 4C,E). To uncover pathways associated with CSF1R independence, we performed KEGG analysis [33] on all upregulated genes in PLX cells and found that translation, metabolic pathways, and phagosome processes were enriched (Figure 4F). This suggested that CSF1R independent microglia may be in an altered metabolic state due to phagolysosomal processes. Altogether, we conclude that CSF1R- independence is linked to increased phagocytosis, lysosomal function, and altered microglia metabolic states, and that homeostatic microglia are more dependent on CSF1R signaling for survival.

### Mer and CR3 are required for apoptotic RGC clearance

To understand the role of apoptotic cell phagocytosis in promoting distinct microglial properties, we sought to test whether recognition receptors were important both for driving microglial remodeling states as well as CSF1R independence. We first wanted to confirm that microglia were important for clearance of dying neurons within the retina and identify the recognition receptors required, focusing on apoptotic RGCs. We performed a screen of genetic KO of receptors previously implicated in the finding, recognition, or phagocytosis of apoptotic cells: find-me pathway, fractalkine receptor CX3CR1 [36], integrin receptor complement receptor 3 (integrin α_M_ß_2_, CD11b, CR3), TAM receptors Mer and Axl (Figure 5A) [25]. We used wildtype animals from the CX3CR1 (CX3CR1-WT) and Mer (Mertk-WT) background as well as CX3CR1-GFP/+ [37] for controls. First, we looked at whether loss of any of these receptors resulted in changes to microglial density. By wholemount immunostaining, we found slight variations at P5 but none that reached significance compared to CX3CR1-GFP/+ (Supplemental Figure 7A,B). Next, we analyzed the density of total apoptotic bodies by cleaved caspase 3 (CC3) in the ganglion cell layer shortly after peak RGC death to measure any buildup or failure to clear (Supplemental Figure 7C,D and Supplemental Figure 8A). We found that loss of CX3CR1, CR3, Mer, and both Mer/Axl resulted in increased apoptotic body density compared to WT controls suggesting these pathways were important for microglial phagocytosis of dying cells within the retina. To ask what pathways were important for clearing dying RGCs specifically, we looked at the density of CC3^+^RBPMS^+^ double-positive cells at P5 in all KOs (Figure 5B,C,D, Supplemental Figure 8B). We found the CR3 KO, Mertk KO, and Mertk/Axl dKO all had increased density of dying RGCs compared to controls (Figure 5B,C) with no change in overall RGC density or retinal blood vessel development (Supplemental Figure 7E,F,G,H). CX3CR1 and Axl were dispensable for clearance of dying RGCs and Mertk/Axl dKOs did not appear to have a further clearance deficit above Mertk KO alone (Figure 5B,C). Thus, both CR3 and Mer receptors, which are broadly expressed in microglia, are important for the timely clearance of RGCs undergoing developmental cell death, consistent with prior studies implicating these pathways in efferocytosis in the CNS.

**Figure 5.**
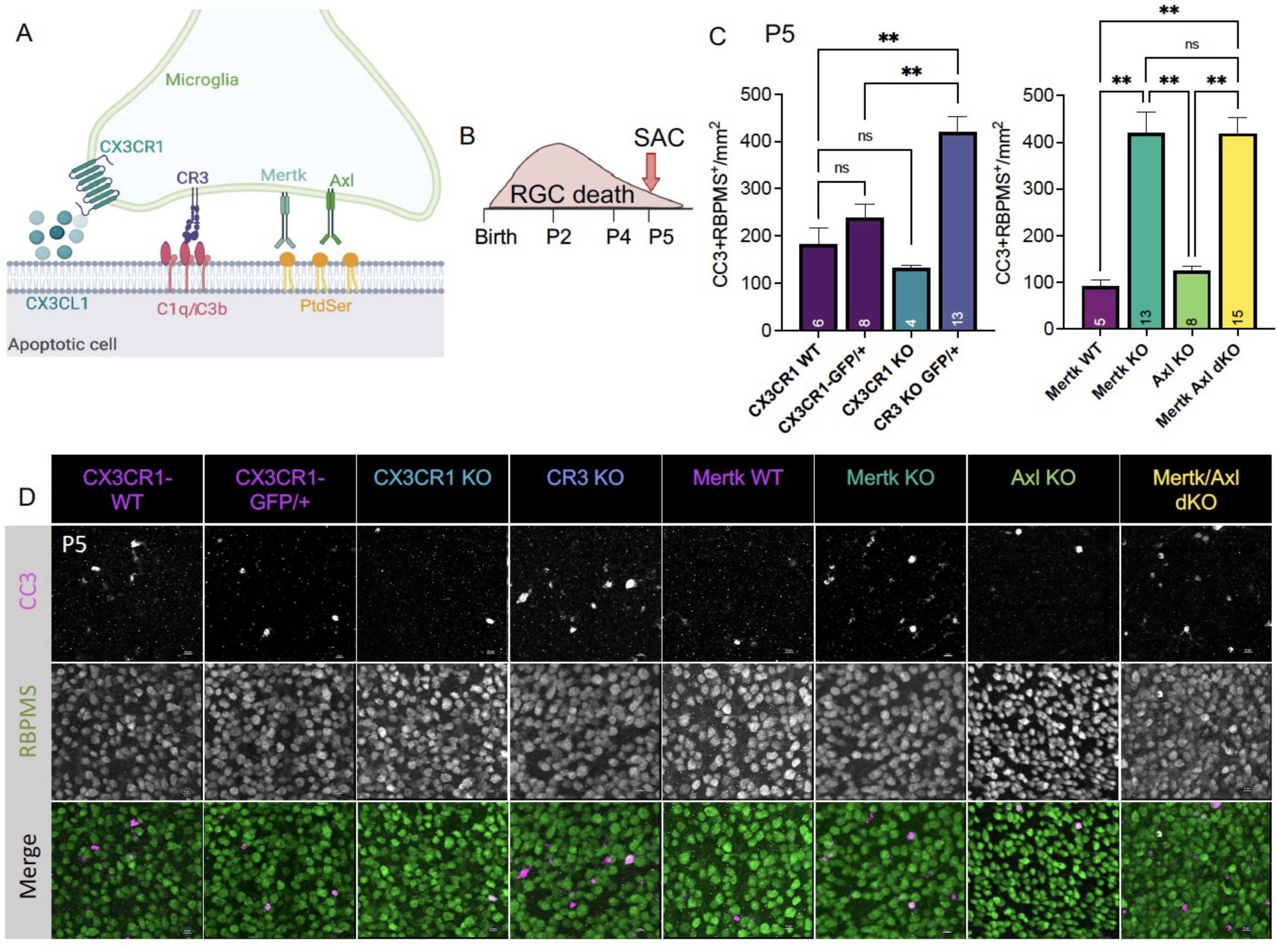
MerTK and CR3 are required for apoptotic RGC clearance. (A) Cartoon schematic of candidate pathways. (B) Schematic for collection at P5 after the bulk of RGC developmental death. (C) Quantification of the average of central and peripheral 0.4 mm^2^ RGC death (±SEM) (Figure S9C) in dorsal leaf of all genotypes (n=6 CX3CR1 WT, n=8 CX3CR1-GFP/+, n=4 CX3CR1 KO, n=13 CR3 KO, CX3CR1-GFP/+, n=5 Mertk WT, n=13 Mertk KO, n=8 Axl KO, n=15 Mertk Axl dKO). (left) Welch’s ANOVA test W(3,12.58)=23.75, ****p<0.0001 and Dunnett’s T3 multiple comparisons test. CX3CR1 WT vs CX3CR1-GFP/+ ns, CX3CR1 WT vs CX3CR1 KO ns, CX3CR1 WT vs CR3 KO **p=0.0016, CX3CR1-GFP/+ vs CX3CR1 KO *p=0.035, CX3CR1-GFP/+ vs CR3 KO **p=0.004, CX3CR1 KO vs CR3 KO ****p<0.0001. (right) Kruskal-Wallis test statistic=25.97 ****p<0.0001 and Dunn’s multiple comparisons test. Mertk WT vs Mertk KO **p=0.001, Mertk WT vs Axl KO ns, Mertk WT vs Mertk Axl dKO **p=0.0014, Mertk KO vs Axl KO **p=0.0026, Mertk KO vs Mertk Axl dKO ns, Axl KO vs Mertk Axl dKO **p=0.0036. Not all comparisons shown on graphs. (D) Max projected confocal images of dying RGCs (CC3^+^RBPMS^+^) in KOs in the dorsal mid-periphery in the ganglion cell layer. Apoptotic bodies, CC3 (magenta); RGCs, RBPMS (green). Scale bars 10µm.

### Axl promotes microglial survival in the absence of CSF1R signaling

Having identified receptors important for RGC clearance (CR3 and Mer), we were next able to ask whether these pathways were also important for driving microglial survival following CSF1R inhibition. We administered PLX3397 as previously published [19] (Figure 6A) and, surprisingly, found that loss of CR3, CX3CR1, or Mer did not significantly reduce the proportion of surviving microglia compared to controls (Figure 6B,C). However, loss of Axl allowed for greater depletion, matching levels achieved with loss of developmental apoptosis (Bax KO) that we previously reported [19] (Figure 6B,C). Mertk/Axl dKOs did not have an additive effect, suggesting Axl signaling alone was important for microglial survival in the absence of CSF1R signaling (Figure 6B,C). Altogether, we find that the receptors important for effective clearance (Mer and CR3) do not alter microglial dependence on CSF1R, but that TAM receptor Axl, which is induced by neuronal apoptosis, augments microglial survival in the absence of CSF1R inhibition in retina.

**Figure 6.**
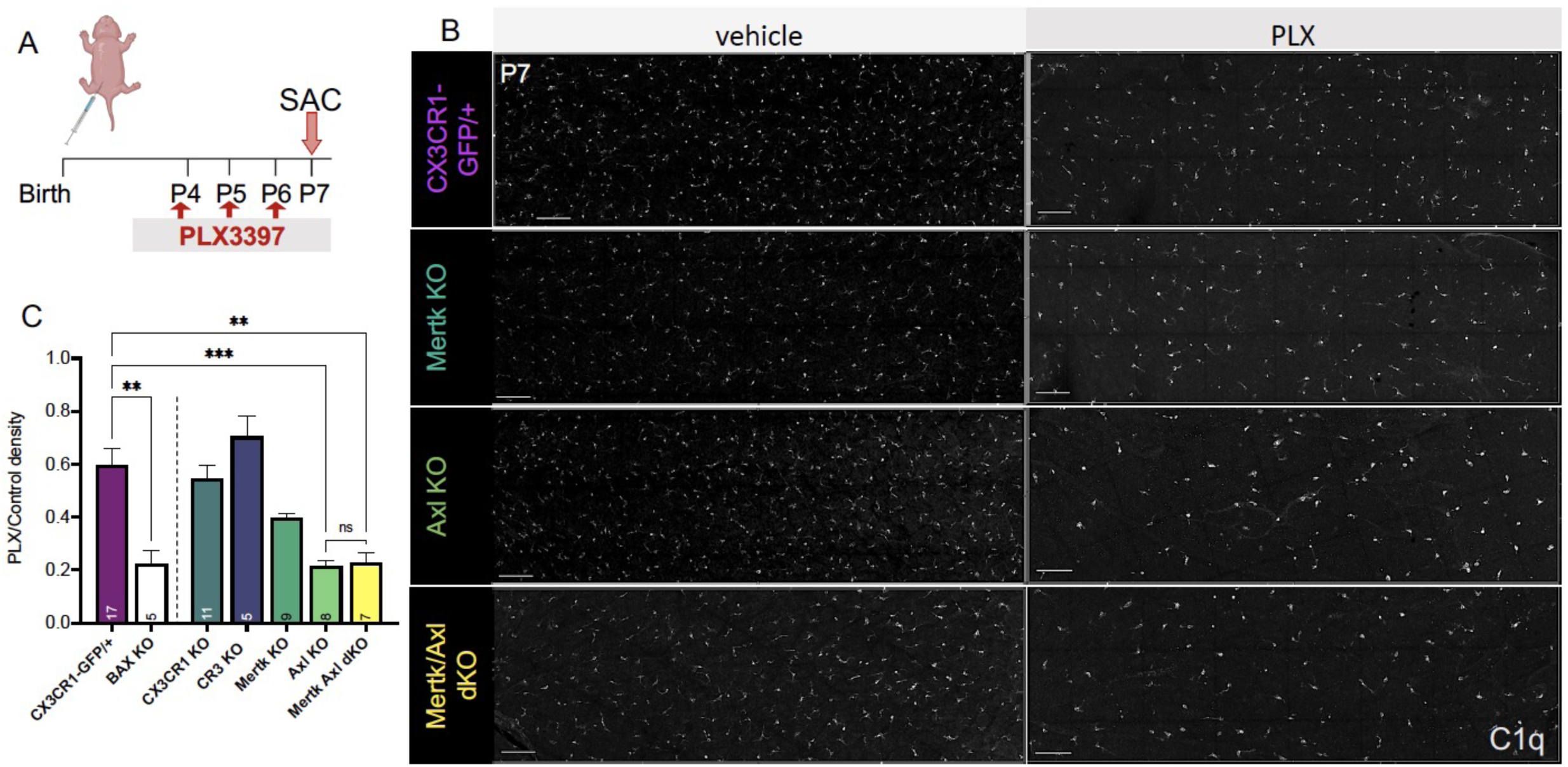
Axl signaling promotes microglial survival in the absence of CSF1R signaling. (A) Dosing regimen of PLX3397 to various genotypes. (B) Confocal images of microglia in the NFL/GCL in all genotypes from central to mid-periphery of the dorsal leaf. C1q (mono). Scale bars 100µm. (C) Ratio of density of CD45^+^CX3CR1-gfp^+^ or CD45^+^CD11b^+^ (microglia/singlets) in PLX treated retinas over genotype-matched controls by flow cytometry. CX3CR1-GFP/+ and BAX KO data was previously published [19]. (n=17 CX3CR1-GFP/+, n=5 BAX KO, n=11 CX3CR1 KO, n=5 CR3 KO CX3CR1-GFP/+, n=9 Mertk KO, n=8 Axl KO, n=7 Mertk Axl dKO; ±SEM). Welch’s ANOVA W(6,18.55)=16.53, ****p<0.0001 and Dunnett’s T3 multiple comparisons test. GFP/+ vs BAX KO **p=0.005, GFP/+ vs CR3 KO ns, GFP/+ vs CX3CR1 KO ns, GFP/+ vs Mertk KO ns, GFP/+ vs AXL KO ***p=0.0004, GFP/+ vs Mertk Axl dKO **p=0.001, BAX KO vs CR3 KO *p=0.0168, BAX KO vs CX3CR1 KO *p=0.0151, BAX KO vs Mertk KO ns, BAX KO vs Axl KO ns, BAX KO vs Mertk Axl dKO ns, CR3 KO vs CX3CR1 KO ns, CR3 KO vs Mertk KO ns, CR3 KO vs AXL KO *p=0.0185, CR3 KO vs Mertk Axl dKO *p=0.0175, CX3CR1 KO vs Mertk KO ns, CX3CR1 KO vs Axl KO ***p=0.0008, CX3CR1 KO vs Mertk Axl dKO **p=0.0021, Mertk KO vs Axl KO ****p<0.0002, Mertk KO vs Mertk Axl dKO *p=0.0394. Axl KO vs Mertk Axl dKO ns. Not all comparisons shown.

### Mer and Axl are not required for expression of lysosomal, lipid metabolism, or remodeling genes

Since Mer was important for phagocytosis while Axl mediated survival in the absence of CSF1R signaling, we wanted to delineate microglial gene expression changes that were driven by Mer versus Axl. We performed bulk RNA-seq on sorted microglia from P4 Mertk and Axl KOs and compared them to WT controls by DESeq2 (Figure 7A). Compared to WT controls, we found modest changes in gene expression, with 42 downregulated genes and 44 upregulated genes in Mertk KO microglia (Figure 7B,D, Supplemental Table 4). However, despite the fact that Mertk KOs had reduced clearance of dying RGCs, there was no effect on lysosomal or lipid metabolism genes. Further, DAM/ATM/PAM-related genes were also not significantly reduced, but, in fact, *Csf1* and *Itgax* were significantly increased (Figure 7B,D).

**Figure 7.**
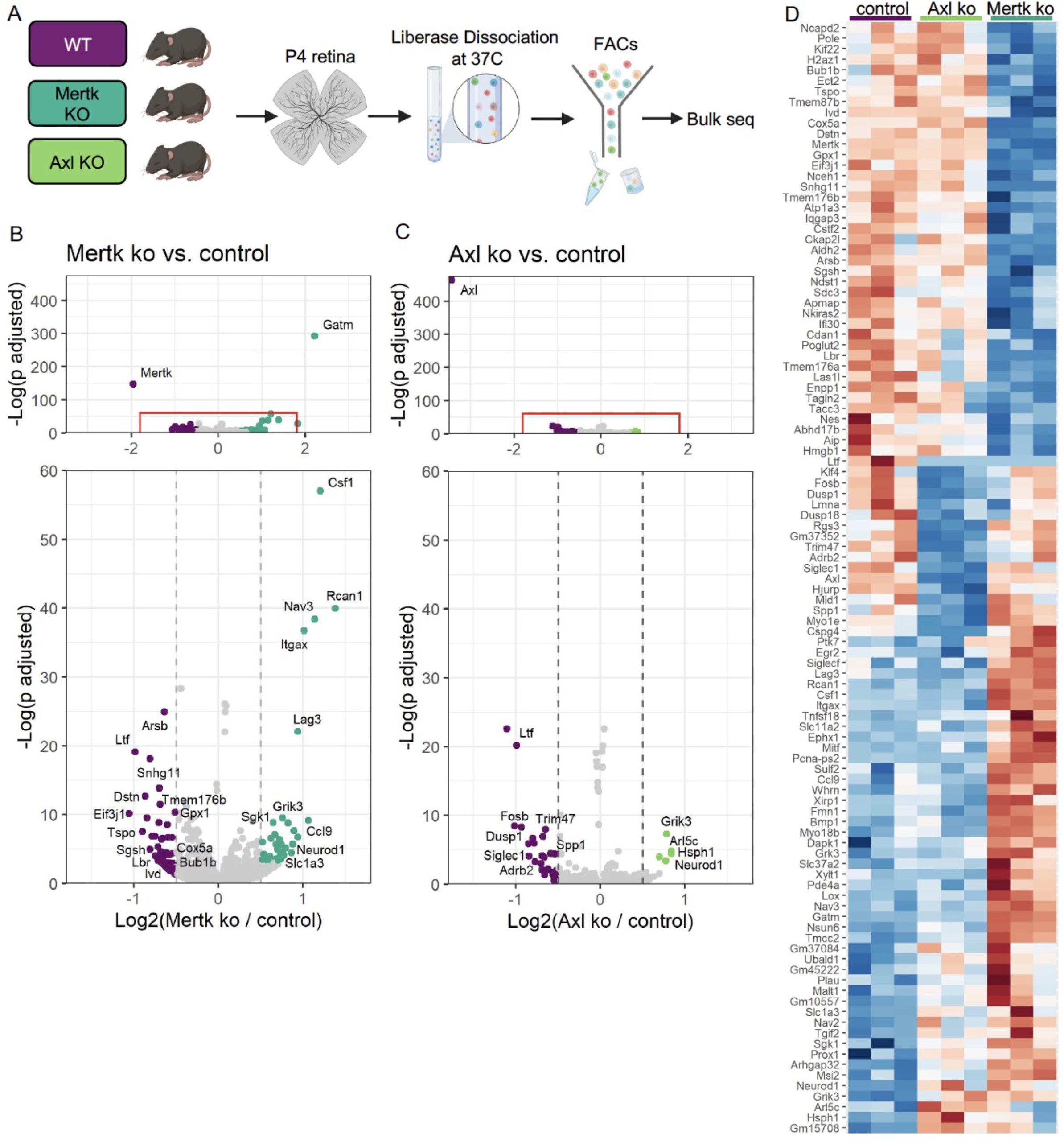
MerTK and Axl are not required for expression of lysosomal, lipid metabolism, or remodeling genes. (A) Workflow for P4 retinal collection, dissociation, sorting, and sequencing of individual microglia from 3 different groups: WT, Mertk KO, and Axl KOs (n=3 each). (B,C) Volcano plot of differentially expressed genes. Top: showing all genes and bottom: zoomed in as indicated in red square. Each gene is plotted according to the significance (-Log(p value)) and magnitude (Log(fold change)) of the difference such that those genes enriched in KO have Log(fold change) > 0. Colored points indicate genes with p-value ≤ 0.05 and Log(fold change) > 0.5. (B) Comparison of Mertk KO relative to WT and (C) comparison of Axl KO relative to WT. (D) Heatmap of all differentially expressed genes between WT compared to Mertk KO and WT compared to Axl KO. Color represents z score of normalized gene expression across all samples. Red=high, blue=low.

Comparison of Axl KO to WT microglia revealed even fewer differentially expressed genes, with no change in lysosomal, lipid metabolism, or DAM/ATM/PAM-related gene expression, except the downregulation of *Spp1*, and no change in genes associated with survival pathways (Figure 7C,D, Supplemental Table 4). The majority of differentially expressed genes were different in the two KOs (Figure 7D); however, sequencing of microglia from P7 Mertk/Axl dKO retinas again revealed no change in lysosomal, lipid metabolism, or DAM/PAM/ATM-related genes (Supplemental 9A,B, Supplemental Table 5). Based upon these findings, we first conclude that while phagocytosis is altered in Mertk KOs, this is not sufficient to suppress expression of remodeling genes, suggesting that Mer-mediated signaling does not strictly drive microglia remodeling gene expression. Furthermore, while Axl facilitates microglia survival after CSF1R inhibition, it is also not required to drive microglia remodeling gene expression, raising the possibility that Axl itself, through recognition of apoptotic neurons, could be mediating survival.

## Discussion

scRNAseq has revealed that microglia in development and disease are transcriptionally diverse across the CNS [38]. Due to their remarkable plasticity, a major challenge is to determine the impact of local cues on microglial state and properties. Here, using the postnatal retina as a model system, we establish neuronal cell death as a key factor driving developmental microglia diversity. We show by scRNAseq that multiple microglial states coexist within this discrete CNS region, encompassing a spectrum from homeostatic to remodeling states with high lysosomal and lipid metabolism gene expression. We establish that multiple distinct remodeling states are driven by neuronal apoptosis, and further show that several of these remodeling states survive CSF1R inhibition, while more homeostatic microglia are susceptible. CR3 and Mer are important for clearance of apoptotic retinal neurons, while Axl facilitates microglial survival following inhibition CSF1R, but neither significantly regulates expression of microglial remodeling genes. Thus, we conclude that cell death is a critical cue for driving diverse microglial properties and that multiple pathways contribute.

We identify several states of remodeling microglia characterized by reduced homeostatic gene expression and elevated lipid metabolism and lysosomal genes, with more discrete expression of various genes including chemokines/cytokines. Many of the genes in these remodeling states are shared with microglia in developing white matter tracts of the brain (CD11c/ATM/PAM) [6, 7, 39] and disease (DAM) [20–22]. Unlike in brain, we find they represent a majority of microglia in the early postnatal retina, with most of them dependent upon neuronal apoptosis. Due to the large proportion of microglia expressing these remodeling genes, we broaden our understanding of microglia subsets present during normal development. For example, we find genes such as *Apoe* and *Ctsd* are widely expressed at varying levels, while *Fabp5* and *Spp1* are confined to more discrete subsets, suggesting that regulation of these genes and microglial states is multifaceted. At present, it remains unclear whether they transition from state to state. Since we find an increase in the proportion of homeostatic microglia in Bax KO retinas, this argues that microglia in the context of neuronal death shift from a homeostatic state to more remodeling ones.

In other contexts, efferocytosis can have a major impact on the transcriptional profile and state of the phagocyte. For example, phagocytosis promotes distinct transcriptional states in macrophages in a tissue-specific manner throughout the body [40], and tissue macrophages modify their repertoire of phagocytic machinery in response to phagocytosis [41]. In the CNS, there is much to learn about how phagocytosis impacts microglial state and function [13, 42]. One study addressing this found engulfment of newborn neurons by microglia in adult neurogenic zones stimulates a microglial secretome that subsequently regulates neurogenesis [43]. Here, in the postnatal retina, phagocytosis and digestion appear to be a major aspect of microglial heterogeneity. We postulate that some remodeling states may even represent distinct stages of the clearance process. Our analysis also identified microglial states that are not regulated by neuronal death, arguing they may be involved in other important microglial functions during postnatal retinal development. For example, astrocytes are also pruned by microglia in the postnatal retina, which has been shown to be Bax-independent and also not require CR3, Mer, or CX3CR1 pathways [44]. Future studies could target these specific microglia subsets and link them to other important remodeling functions in the developing retina.

Despite the fact that postnatal RGC death has been a well-characterized model of developmental cell death of the CNS [16], the recognition pathways required for the clearance of apoptotic RGCs was previously undefined. CR3 and Mer are well-known recognition receptors in the clearance of stressed or dying neurons in various contexts [25]. Here, we show that both receptors are important for clearance of apoptotic RGCs. Previous work has suggested that complement component C1q may get deposited on exposed PtdSer [45], suggesting the pathways might work together to signal for phagocytosis, but whether complement and TAM pathways are working redundantly or cooperatively is unknown in any context. While receptors Mer and Axl are both members of the TAM receptor family of tyrosine kinases [46], it has been appreciated that Mertk is more stably expressed in microglia, while Axl is more dynamic and upregulated with inflammation or disease [47]. Both have previously been implicated in phagocytosis and in regulating inflammatory responses [48], however this is cell type-dependent [49]. Here, we find Mertk/Axl dKO did not have an enhanced clearance deficit for apoptotic RGCs compared to Mertk KO, although this could be due to a plateau effect since RBPMS likely gets downregulated as apoptotic programs continue [50].

Although Mer is important for clearance, we did not see a dramatic reduction in lysosomal or lipid metabolism genes in either Mertk or Axl KO microglia. These results are consistent with disease contexts where loss of Mertk and Axl do not have a dramatic effect on the DAM signature in mouse model of Alzheimer’s disease [51]. This suggests that transcriptional changes in postnatal retinal microglia may depend upon other pathways for apoptotic cell recognition. An alternate possibility is that Mertk KO microglia are still clearing, but at a reduced rate, since live-imaging has shown that loss of Mer results in delayed engagement with apoptotic cells but does not abolish phagocytosis completely [52].

We previously found that a large proportion of microglia in the postnatal retina were more resistant to inhibition of the important survival pathway CSF1R [19]. Here, we identify specific remodeling states of microglia that persist following CSF1R inhibition showing reduced homeostatic gene expression. Other groups recently found small populations of CSF1R-independent or repopulating microglia in the brain, and we note similar patterns of gene expression, with reduced homeostatic genes such as *Tmem119* and *P2ry12*, and increased expression of genes such as *Lyz2* [34, 53]. In these microglia repopulation studies, it was proposed that more immature microglia survive CSF1R inhibition, but microglia maturation may not be a key factor in the context of development since we previously found greater dependence of retinal microglia on CSF1R at embryonic stages than in postnatal retina [19]. We find that most of these resistant microglia subsets are induced by neuronal apoptosis, consistent with our prior analysis showing increased microglial dependence on CSF1R in Bax KO retina [19]. Interestingly, in culture, peritoneal and bone marrow macrophages that phagocytose apoptotic cells also survive in the absence of serum or survival factors [54]. In periods of elevated cell death and possible lack of extrinsic survival factors, such as neuronally expressed CSF1R ligand, IL-34 [55], phagocytes such as microglia might have evolved strategies to survive in order to continue their job of maintaining homeostasis or reducing inflammation [54]. Importantly, cholesterol, along with CSF-1/IL-34 and TGF-ß2, has been shown to be an important survival factor for microglia in culture [56]. We observed that PLX-treatment resulted in enrichment of microglia with the lowest levels of homeostatic gene expression and high levels of lysosomal and lipid metabolism genes. Thus, CSF1R signaling may also be important for promoting microglial homeostasis [57]. Consistent with this, mutations in CSF1R can lead to leukoencephalopathy, a neurodegenerative disorder which is associated with loss of homeostatic microglia phenotype [35]. It will be valuable for future studies to explore the contributions of CSF1R signaling to microglial survival and homeostasis during development.

Here, we find that Axl expression, which is regulated by neuronal apoptosis, enhances microglial survival in the absence of CSF1R signaling. Since loss of Axl did not substantially alter expression of other candidate apoptotic or survival pathways in microglia, one possibility is that signaling directly downstream of Axl is involved. Axl has been shown to inhibit apoptosis in a variety of cell types [58] including fibroblasts [59], oligodendrocytes in contexts of growth factor withdrawal or TNF toxicity [60], and in Gonadotropin-releasing hormone neurons in the brain [61]. Its role in various types of cancer has been well-established for promoting tumor growth and metastasis in part due to its ability to inhibit apoptosis of cancer cells [62]. Axl has diverse downstream signaling cascades depending on context [58], but one candidate signaling cascade is the PI3K-Akt-NfkB-Bcl2 pathway. Activation of PI3K and Akt has shown to be important for survival of cultured macrophages [54], oligodendrocytes [60, 63], neurons [61] as well as cancer cells [58].

Altogether, we show neuronal cell death is a key factor driving multiple states of microglia in developing retina, with distinct transcriptional profiles and independence from CSF1R signaling for survival. Apoptotic cell recognition by microglia is thus a critical developmental event driving diverse effects on gene expression, clearance and survival, with distinct pathways involved in mediating each of these responses.

## Acknowledgements

This research was supported by the National Eye Institute and the National Institute of Neurological Disorders and Stroke of the National Institutes of Health under Awards R01EY030307 (MLV), and T32EY024234, T32NS115664 (NG). We thank the University of Utah BPRB Animal Facility, University of Utah Flow Cytometry Core, University of Utah High Throughput Genomics Core, and University of Utah Bioinformatics Core for technical support.

## Materials and Methods

### Experimental Model and Subject Details

#### Animal husbandry and procedures

All animals were treated within the guidelines of the University of Utah Institutional Animal Care and Use Committee (IACUC) and all experiments were IACUC approved. Mice were housed in an AAALAC accredited animal facility with 12h light/12h dark cycles and ad libitum access to food and water. Both sexes were used for all experiments. Information on the ages of mice used for each experiment can be found in the figures/text. Pexidartinib (PLX3397) (AdooQ BioScience A15520) was dissolved in corn oil (Sigma-Aldrich C8267) and 10% DMSO (Fisher Scientific BP231) and administered to postnatal pups by daily intraperitoneal injection P3-P5 or P4-P6 at 0.25 mg/g body weight. Mice were euthanized by isoflurane asphyxiation followed by decapitation.

#### Mouse strains

The B6.CX3CR1-GFP/+ mice were a gift from Richard Lang with permission from Dr. Steffen Jung [37]. Bax KO mice (JAX 002994) and B6. CD11b KO (JAX 003991) were purchased from the Jackson Laboratory. The B6.Mertk KO and B6.Axl KO strains were a kind gift from Dr. Greg Lemke.

### Method Details

#### Tissue processing

Following euthanasia, retinas were dissected in ice-cold 0.1M PBS. Whole heads were fixed in 4% PFA for 45 minutes to an hour. Heads were washed 3 times for 15 minutes in PBS and underwent 12-16-hour consecutive treatments with 15% and 30% sucrose in PBS at 4°C. Heads were then embedded in OCT compound (Ted Pella 27050), stored at -80°C, and sectioned at 16μm thickness. For retinal whole mounts, eyes were removed from the head and retinas were carefully dissected from the rest of the eye (cornea, lens, RPE, hyaloid vasculature, vitreous, ciliary body) in ice-cold PBS. Whole neural retinas were washed in PBS for 10-20 minutes and then fixed in 4% PFA for 30-45 minutes at room temperature. For RNAase-free dissections for qHCR or FACs, retinas were carefully dissected in RNase-free conditions using ice-cold, sterile RNase-free PBS, removing all non-neural eye tissue (ciliary body, pigmented epithelium, vitreous).

#### Immunohistochemistry

As previously done [19], frozen sections were incubated at 37°C for 30 minutes, post-fixed for 15 minutes in 4% PFA and washed in ice cold PBS for 3 times for 5 minutes. Then they were blocked for 1 hour at room temperature (0.2% triton-X, 10% BSA, 10% normal donkey serum in 0.01M PBS), and incubated in primary antibody overnight at 4°C in (0.2% triton-X, 5% BSA in 0.01M PBS). The following day, sections were washed 3 times with PBS and incubated in secondary antibodies (5% BSA in PBS) for 2 hours at room temperature, washed, and mounted with Fluoroshield mounting medium with DAPI (MilliporeSigma F6057). For whole mount immunostaining, retinas were incubated in primary antibody at 4°C for 2 days. Primary antibodies used: goat GFP (abcam ab5450), rabbit C1q 1:1500 (abcam ab182451), rabbit active caspase-3 1:500 (CC3) (BD Biosciences 559565), guinea pig RBPMS 1:1000 (MilliporeSigma ABN1376MI), FITC IB4-lectin 1:400 (Sigma L9381). Secondary antibodies: 488 Donkey anti-goat 1:400 (Invitrogen A11055), 555 Donkey anti-rabbit (Life Technologies A31572), and 647 Donkey anti-guinea pig (Jackson ImmunoResearch 706-605-148).

#### In situ hybridization chain reaction (HCR)

Wholemount retinas were fixed overnight in 4% PFA in 4°C. Retinas were washed and dehydrated in methanol/PBS at 25%, 50%, 2 times 100% for 15 minutes each, stored in 100% methanol overnight at 4°C and 20°C long term. *In situ* hybridization was performed as published [64] using v3.0 reagents from Molecular Instruments (https://www.molecularinstruments.com). Briefly, samples were rehydrated using (75% Methanol/25% PBST, 50% Methanol/50% PBST, 25%, Methanol/75% PBST, 2 times 100% PBST), treated with Proteinase K, and post fixed 20 minutes at room temperature in 4% PFA, and washed 3 times with PBST. Pre-hybridization was performed in 30% probe hybridization buffer for 30 minutes at 37°C, and retinas were placed in hybridization buffer at 37°C overnight. Retinas were washed, placed in amplification buffer for 30 minutes at room temperature. Separately, hairpins used for amplification were denatured at 95°C for 90 seconds and cooled to room temperature for 30 minutes. Retinas were placed in amplification buffer with hairpins at room temperature in the dark overnight. Retinas were washed, DAPI stained, and mounted on slides. Probes recognizing all known transcript variants for each of Cx3cr1 and Ccl3 were generated by Molecular Instruments. For Cx3cr1-GFP knockin mice, the fluorescent signal represents detection of Cx3cr1 mRNA by HCR and GFP fluorescence which persists through the procedure.

#### Confocal Microscopy

Confocal images were acquired on an inverted Nikon A1R Confocal Microscope. Images were acquired at 20X objective with a 3X digital zoom. Multi-points were stitched with a 10% overlap. Images of retinal whole mounts were 144 multi-point images (on average) to obtain the entire dorsal retina. Stacks through the Z plane were at 0.8µm steps of about ∼13 µm thickness to capture just the nerve fiber layer (NFL) and ganglion cell layer (GCL) at 0.2μm pixel resolution. Whole mount retina images represent max projections of inner retina (NFL/GCL) from the central retina (optic nerve) to the periphery (edge of retina). In cases of microglial quantification, whole mount imaging and analysis spanned ∼25µm thickness from NFL to inner plexiform layer (IPL). Image acquisition settings were consistent across ages and genotypes.

#### Fluorescence-Activated Cell Sorting (FACS) and Flow Cytometry

Except for SC-sequencing, we pooled 2 retinas from an individual animal for each sample for flow cytometry and FACs. Freshly dissected pure retinas were dissociated in PBS, 50 mM HEPES, 0.05mg/ml DNase I (Sigma D4513), and 0.025 mg/ml Liberase (Sigma 5401119001) for 35 minutes with intermediate trituration at 37°C. Cells were passed through a 70 µm nylon cell strainer, washed with ice-cold staining buffer (1X PBS, 2% BSA, 0.1% sodium azide, 0.05% EDTA), and red blood cells were lysed (eBioscience 00-4333-57). Cell counts were determined using a cell counter (Invitrogen Countess) and Fc block (BD Biosciences 553142) was added at 2 μL per 10^6^ cells. Antibodies were applied for 30 min on ice. Antibodies used: BV421 CD45 1:200 (BD Bioscience 563890), 488 CD11b 1:200 (CD45 (BD Bioscience 557672), PE CCR2 1:200 (R&D Systems FAB5538P), and APC Ly6C 1:200 (BD Bioscience 560595). Cells were washed, pelleted, and resuspended in 500 μL staining buffer. FACS was performed using a BD FACS Aria cell sorter at the University of Utah Flow Cytometry Core. Forward and side scatter were used to eliminate debris, and both the width and area of the forward and side scatter was used to discriminate singlets. For flow analysis, roughly 1 million singlet events (300,000 in rare cases) were recorded for flow analysis using FlowJo software (Flowjo, LLC, Ashland, Oregon).

#### Single-cell RNA sequencing

13 animals/26 retinas were pooled for each of the Bax WT and littermate KO samples. 12 animals/24 retinas for PLX3397 CX3CR1-GFP/+ and 11 animals/22 retinas were pooled for Vehicle CX3CR1-GFP/+ controls. FACs protocol was performed as normal, except pooling 6 retinas per tube during dissociation. CD45^+^ CD11b^+^/GFP^+^ CCR2^-^ cells were sorted for Bax samples and CD45^+^GFP^+^ Ly6C^-^ cells were sorted for PLX and Vehicle samples. Cells were sorted using a 5 laser BD FACSAria with a 70μm nozzle at 55psi by the University of Utah Flow Core into a cold, empty tube before library generation. 15,000 cells for each Bax sample and 7,000-10,000 cells for PLX and Vehicle were loaded onto 10X chip. Single-cell libraries were generated with 10X Genomics Single Cell 3’ Gene Expression Library Prep v3 reagents at the University of Utah High Throughput Genomics core. The libraries were sequenced on a NovaSeq 6000 to generate at least 200 million paired end reads of 150 bp per sample.

#### Analysis of Single-cell RNA-seq Data

Aligned reads to the mm10 reference from 10X genomics (version 3.0.0 from Ensembl 93) and generated feature-barcode matrices using cellranger count v3.1.0 with expected-cells set to 5000. Filtered feature matrices from cellranger were further filtered with Seurat (v3.1.5) [65]. High-quality cells were selected by < 10% mitochondrial gene content, and high number of features (genes). Feature cut-off was specific to each sample: > 1,000 for Bax-ko, > 1,500 for Bax-wt, > 1,800 for vehicle, and > 2,500 for PLX-treated. Likely doublets, with high transcript counts, were also eliminated: < 25,000 counts for Bax-ko, < 27,000 for Bax-wt, < 47,000 for vehicle, and < 125,000 for PLX treated. In silico genotyping, as described below, was used to assign cells to the correct groups. Filtered cells from all samples were then merged and processed with the SCTransform pipeline [66]. Mitochondrial percentage was regressed out of the model and Bax, Ftl1, and Ftl1-ps1 were excluded from the variable features list used for dimensionality reduction. From here we subsetted the data to include only microglial cells. We identified non-microglia populations by their expression of Ptprc, Plac8, Clec12a, Ms4a7, Mrc1, and Rorb, and reran the SCTransform pipeline as before with the exception that FindClusters was run with resolution set to 0.5. Differentially expressed genes for cluster comparisons and whole-sample comparisons were identified with the FindMarkers function, using default parameters.

#### In Silico Genotyping

During single-cell library preparation, a portion of Bax-wt and Bax-ko cells were allocated to the wrong sample. We used the scSplit pipeline [67] to perform in silico genotyping and reassign these cells to the correct sample. Beginning with the bam file for each sample generated by cellranger, we used samtools view v1.8 [68] to exclude reads with mapping quality < 10, or flagged as unmapped, not primary alignment, fails quality checks, PCR or optical duplicate, or supplementary alignment. These filtered bam files were used to identify single nucleotide polymorphisms (SNPs) with freebayes v1.3.1(https://arxiv.org/abs/1207.3907v2). Indels, MNPs, and complex alleles were filtered from the input to the algorithm. Region was set to chromosome 7, where Bax is located, minimum allele count was set to 2, use-best-n-alleles was set to 2, and minimum base quality was set to 1. Vcffilter v1.0.1 [69] was used to exclude SNPs with quality score <=30. We found that some crucial SNPs were identified in one sample and not the other, so Bcftools merge v1.9 [68] was used to combine all SNPs into a single file. ScSplit count v1.0.0 [67] was used to the combined SNPs to generate count matrices for the filtered cell barcodes identified by cellranger, and scSplit run, with expected number of mixed samples set to 1, genotyped the individual cells based on these matrices. We assigned Bax-ko and Bax-wt genotypes to the cell populations based on Bax, Ftl1, and Ftl1-ps1 expression.

#### Bulk RNA sequencing

For P4 WT, Mertk KO, and Axl KO samples, CD11b^+^ CD45^+^ Ly6c^-^ cells from one animal (2 retinas) were sorted directly into RLT buffer using a 4 laser BD FACSAria at the University of Utah Flow Core and stored at -80°C. 2 samples were pooled prior to RNA isolation so that each replicate was cells from 2 animals/4 retinas. WT samples had 5470, 6134, 7029 cells, Mertk KO samples had 4245, 4038, and 8591 cells, and AXL KO samples had 5790, 5282, and 5071 cells. For P7 WT and Mertk/Axl dKO, CD11b^+^ CD45^+^ cells from one animal (2 retinas) were sorted directly into RLT buffer (QIAGEN 79216) and stored at -80°C. P7 samples were not pooled and thus represent 1 animal/2 retinas. For WT samples, 3783, 4083, and 4982 cells were collected. For Mertk/Axl dKO samples, 2408, 2633, and 2388 cells were collected. RNA from all samples was purified using the RNeasy Plus Micro kit (QIAGEN 74034). Total RNA was hybridized with the NEBNext rRNA Depletion Solution human/mouse/rat (E6310L) to substantially diminish cytoplasmic and mitochondrial rRNA from the samples. Stranded RNA sequencing libraries were prepared as described using the NEBNext Ultra II RNA Library Prep Kit for Illumina (E7770L). Purified libraries were qualified on an Agilent Technologies 2200 TapeStation using a D1000 ScreenTape assay (cat# 5067-5582 and 5067-5583). The molarity of adapter-modified molecules was defined by quantitative PCR using the Kapa Biosystems Kapa Library Quant Kit (cat#KK4824). Individual libraries were normalized to 10 nM, and equal volumes were pooled in preparation for Illumina sequence analysis. Sequencing libraries were chemically denatured and applied to an Illumina NovaSeq flow cell using the NovaSeq XP workflow (20043131). Following transfer of the flowcell to an Illumina NovaSeq 6000 instrument, a 150 x 150 cycle paired end sequence run was performed using a NovaSeq 6000 S4 reagent Kit v1.5 (20028312). Samples were sequenced to a depth of 39-76 million reads.

#### Analysis of Bulk RNA-seq Data

Optical duplicates were removed with clumpify using BBmap v38.34 (https://jgi.doe.gov/data-and-tools/bbtools/bb-tools-user-guide/bbmap-guide/) and default settings, then Illumina adapters were trimmed with cutadapt v1.16 [70] using a minimum overlap of 6 and minimum length of 20. Reference database was generated by STAR v2.7.9a [71] using mouse Ensembl release 104 with overhang set to 124. Trimmed reads were aligned in two pass mode to generate a BAM file, sorted by coordinates. Reads were assigned to the target with the largest overlap, and uniquely aligned, reversely stranded reads were counted with featureCounts v1.6.3 [72]. Differentially expressed genes were identified from counts using DESeq2 v1.32.0 [73]. Features with fewer than 5 reads in every sample were eliminated before DESeq was run. rlog-transformed values were used for sample visualizations.

### Quantification and Statistical Analysis

#### Image analysis

All counts were blinded and manual, performed using Nikon Elements software (Melville, NY). For double-positive CC3^+^RBPMS^+^ or single RBPMS^+^ counts of retinal whole mounts, two ROIs of roughly 0.4 mm^2^ of central and periphery of dorsal retina were analyzed. Images were max projected and roughly 13µm thick, spanning the NFL to the GCL. For CC3^+^ density, the entire dorsal leaf was analyzed, roughly 2-3 mm^2^ again spanning the NFL to GCL only. See also Figure S9. For microglial quantification, whole mount imaging and analysis spanned ∼25µm thickness from NFL to IPL and 0.5625 mm^2^ of the central to mid-peripheral, vascularized retina of the dorsal leaf.

#### Statistical Methods

Detailed statistical information can be found in the figure legends, including tests used, sample size, and precision measures. All image and flow data were analyzed using Prism 9 software (GraphPad, La Jolla, CA). All data were first tested for normality using four tests: Anderson-Darling, D’Agostino & Pearson, Shapiro-Wilk, and Kolmogorov-Smirnov test. If any one test failed, non-parametric tests were used: Mann-Whitney for pair-wise comparisons and Kruskal-Wallis for comparisons of more than three samples. For all t-tests, we applied the Welch’s correction if the two samples had unequal variances determined by an F-test. We tested for heteroscedasticity in groups of 3 or more by a Brown-Forsythe test. If not significantly different, we ran an Ordinary one-way ANOVA with post-hoc Tukey’s multiple comparison test. If the standard deviations were significantly different between groups, we ran a Welch’s ANOVA with post-hoc Dunnett’s T3 multiple comparison’s test. For all data that is presented as the mean, error bars indicate the standard error of the mean, SEM. We used a 95% confidence interval and a p-value of < 0.05 for rejecting the null hypothesis.

### Data and Software Availability

The sequence data reported in this publication have been deposited in NCBI’s Gene Expression Omnibus (GEO).

---

**Figure S1.**
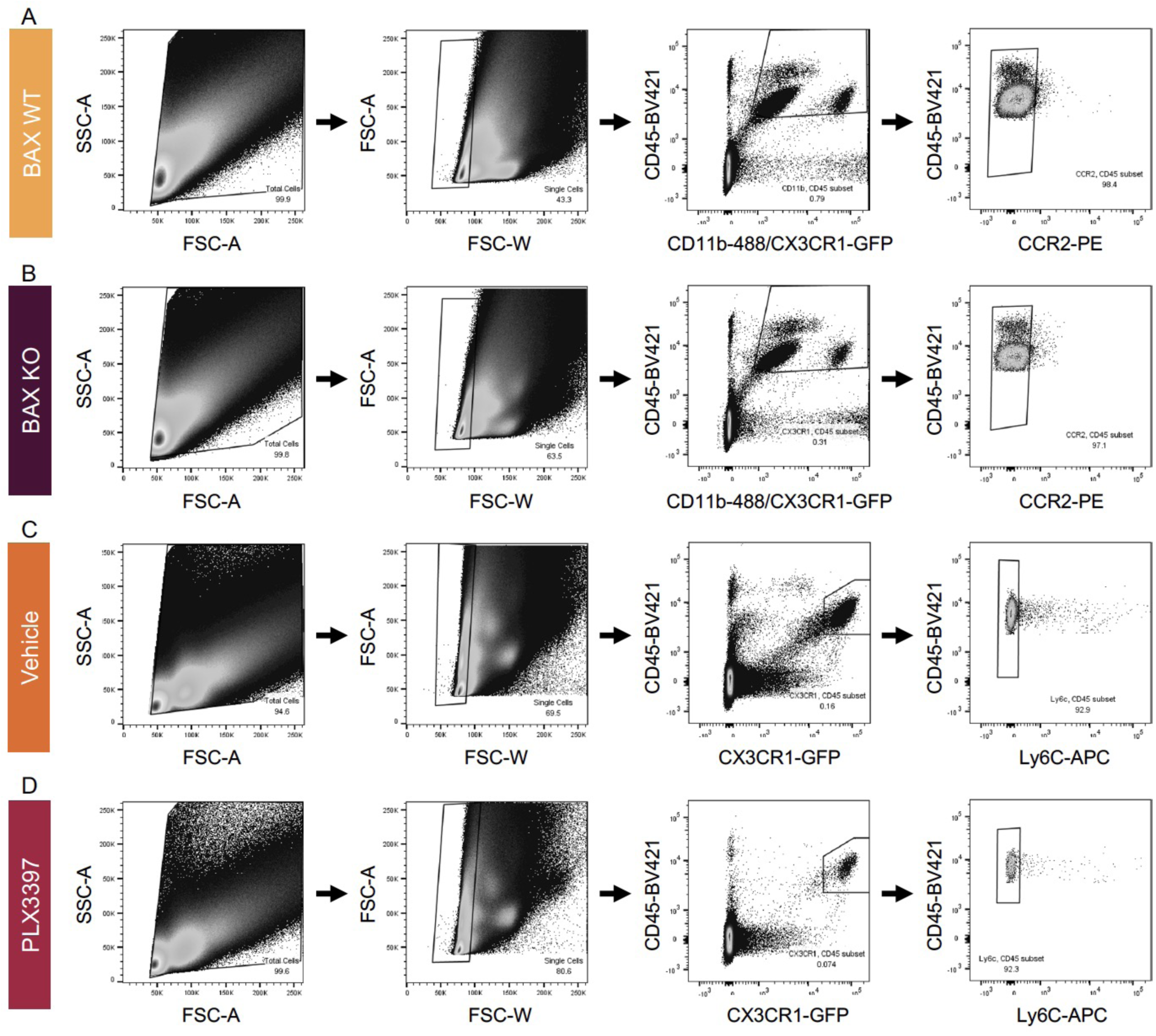
Gating strategy to FAC-sort retinal microglia for single-cell sequencing Related to Figure 1. (A) Gating strategy for FACs of BAX WT and (B) KO samples for single-cell sequencing. Animals were a mixture of CX3CR1-GFP/+ and CX3CR1-+/+. CCR2 was used to exclude monocytes and macrophages. (C) Gating strategy for FACs of CX3CR1-GFP/+ Vehicle and (D) PLX3397 treated samples. Ly6C was used to exclude monocytes and macrophages.

**Figure S2.**
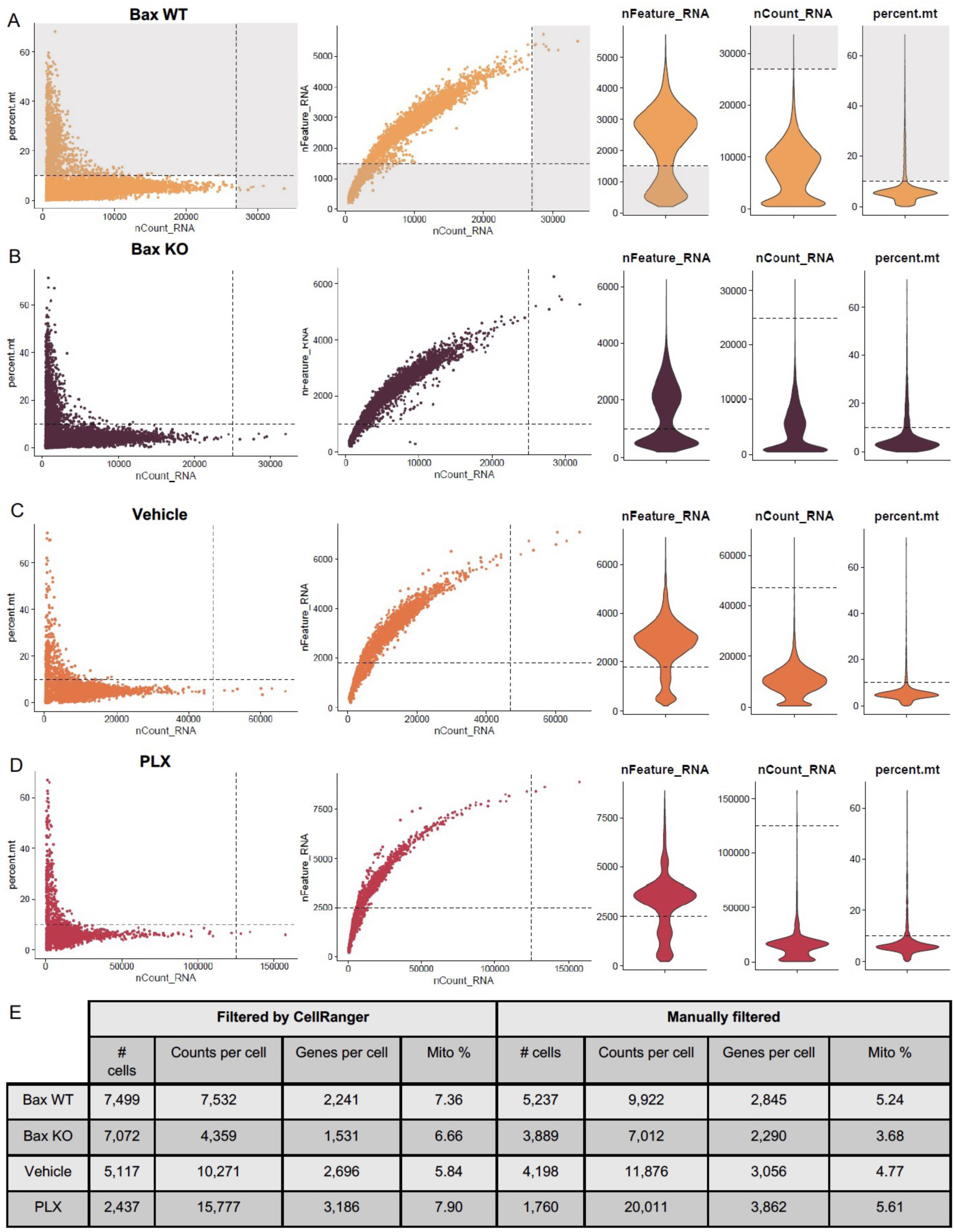
Selection of high-quality cells Related to Figure 1. (A-D) Cells, as identified by CellRanger, were filtered to exclude low quality cells: those with few features (genes) or high mitochondrial content. Cells with especially high transcript counts were eliminated as potential doublets. Cut-offs were set independently for Bax WT (A), Bax KO (B), Vehicle (C), and PLX (D) samples. (E) Dataset composition before and after cell filtering.

**Figure S3.**
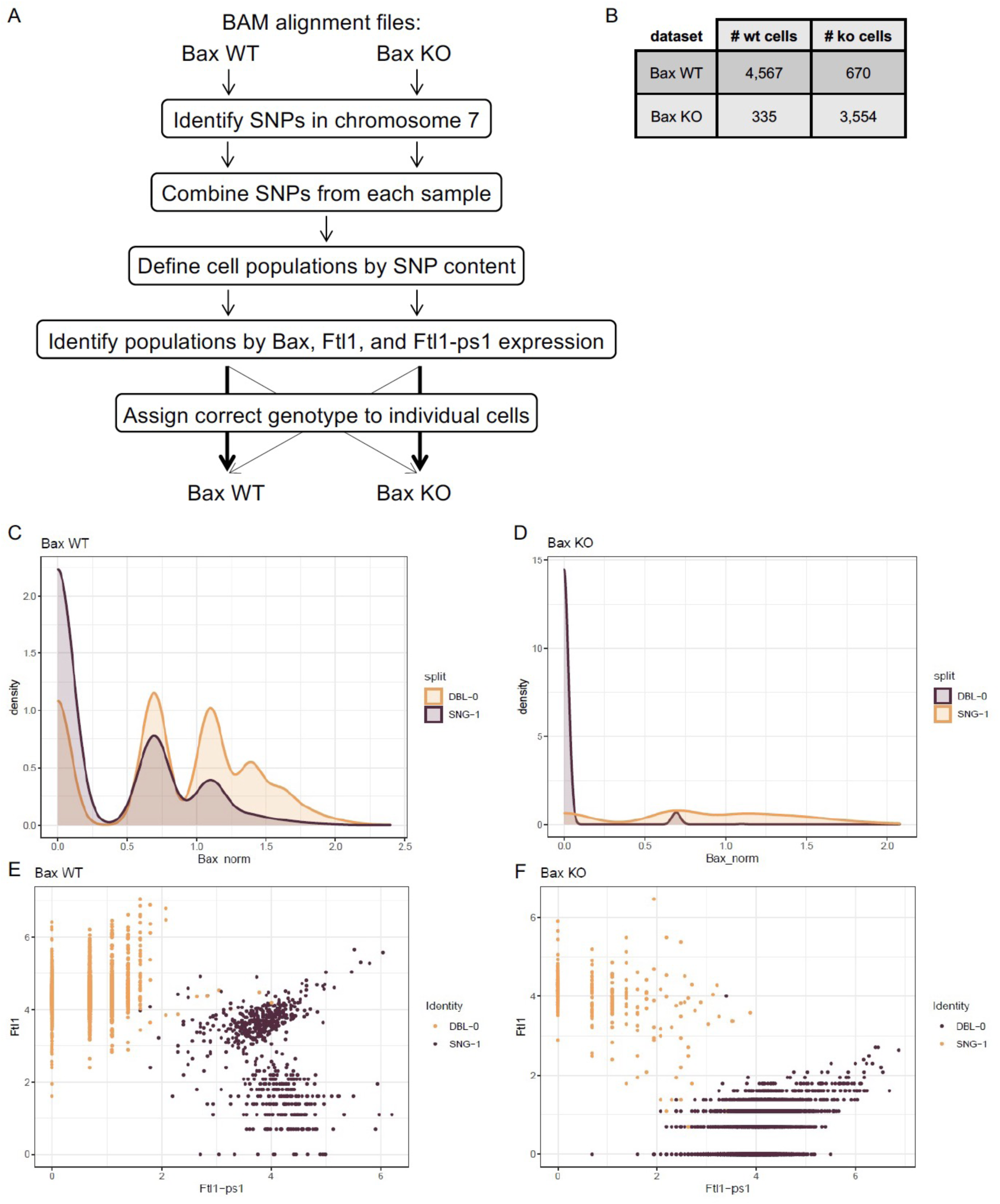
In silico *Bax* genotyping Related to Figure 1. (A) Summary of *in silico* genotyping workflow from alignment files to genotyping of individual cells. (B) Number of cells from the filtered dataset (Bax WT or Bax KO) ultimately assigned to each genotype (# wt or # ko). (C-F) In silico genotyping identified two populations within each dataset, which were classified as wt or ko based on (C, D) Bax, (E, F) Ftl1, and Ftl1-ps1 expression. Data shown are log normalized counts. The two populations are pseudocolored here: yellow for wt and purple for ko.

**Figure S4.**
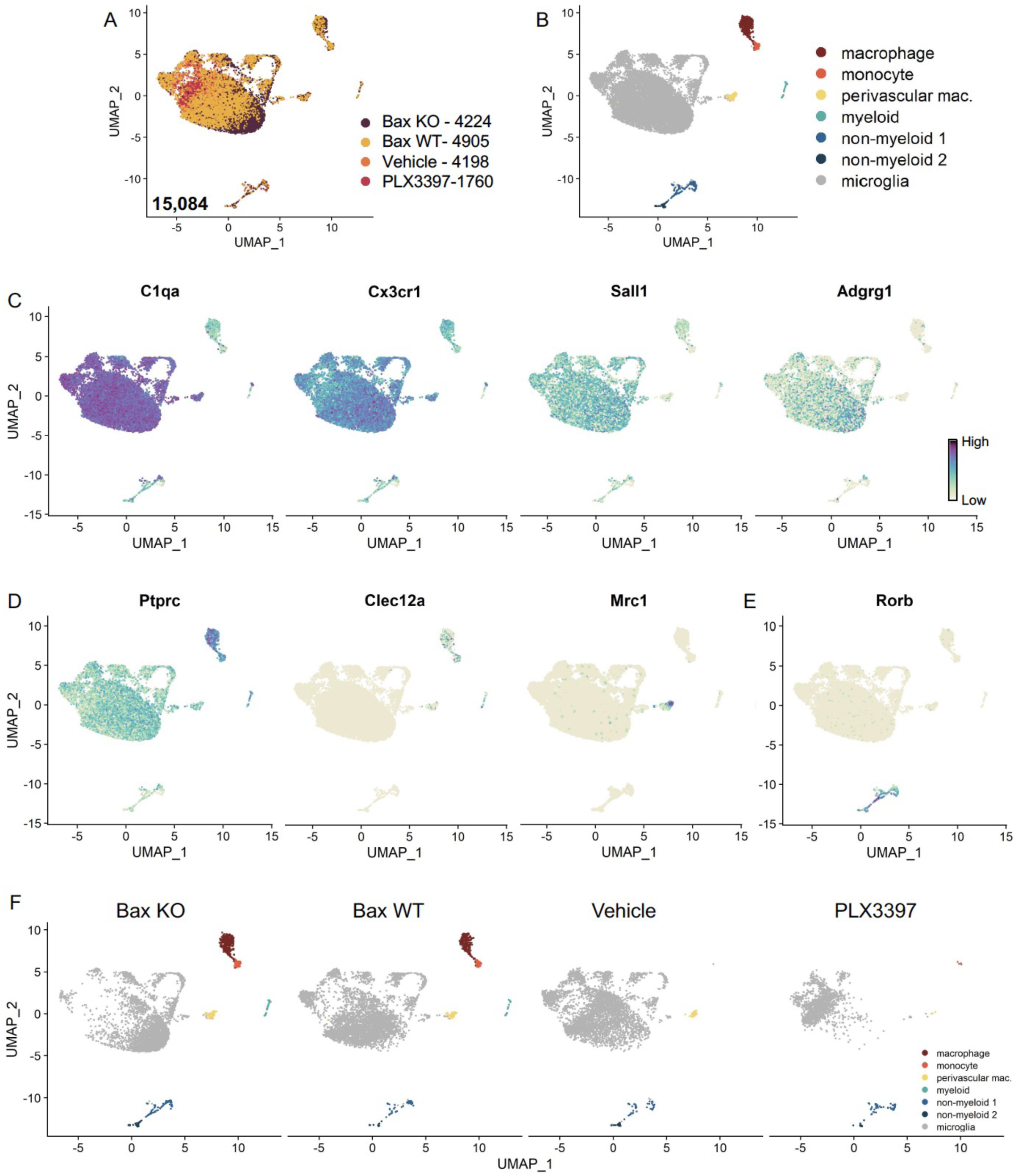
Identification of non-microglia populations Related to Figure 1. (A) UMAP plot of 15,084 cells from all 4 samples. BAX KO, 4224 cells; BAX WT 4902 cells; Vehicle, 4198 cells; PLX3397, 1760 cells. (B) UMAP plot with clusters labeled based on published markers: macrophages (dark red), monocytes (bright red), perivascular macrophages (yellow), other myeloid-like cells (teal), non-myeloid 1 and 2 (shades of blue), and microglia (gray). (C) UMAP plots colored by relative gene expression (dark purple = highest, light yellow= lowest) of highly expressed or specific microglia genes. (D) UMAP plots of published monocyte or macrophage genes. (E) UMAP plot of neuronal gene expression. (F) UMAP plots illustrating the distribution of cells across clusters for each sample.

**Figure S5.**
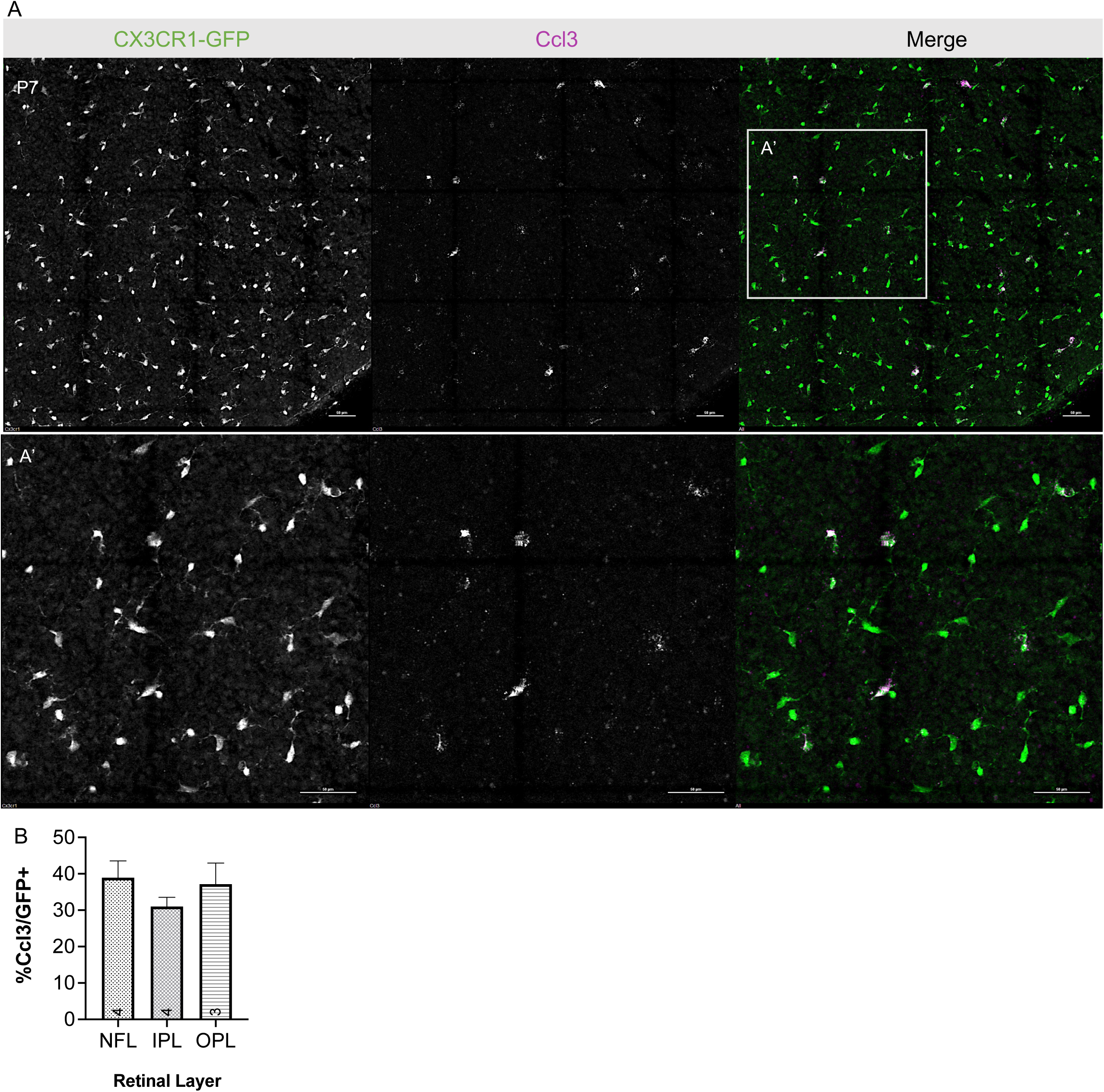
Postnatal retinal microglia express *Ccl3 in vivo* Related to Figure 1. (A) Max projected confocal image of *in situ* hybridization chain reaction (HCR) on P7 whole mount retina probed for *Ccl3*. CX3CR1-GFP (protein, green), Ccl3 (RNA, magenta). A’ inset, higher magnification. Scale bar 50 µm (B) Percent *Ccl3*^+^ of CX3CR1-GFP^+^ cells in each layer of the P7 retina of the mid-periphery. (n=>3; ±SEM). NFL, nerve fiber layer; IPL, inner plexiform layer; OPL, outer plexiform layer.

**Figure S6.**
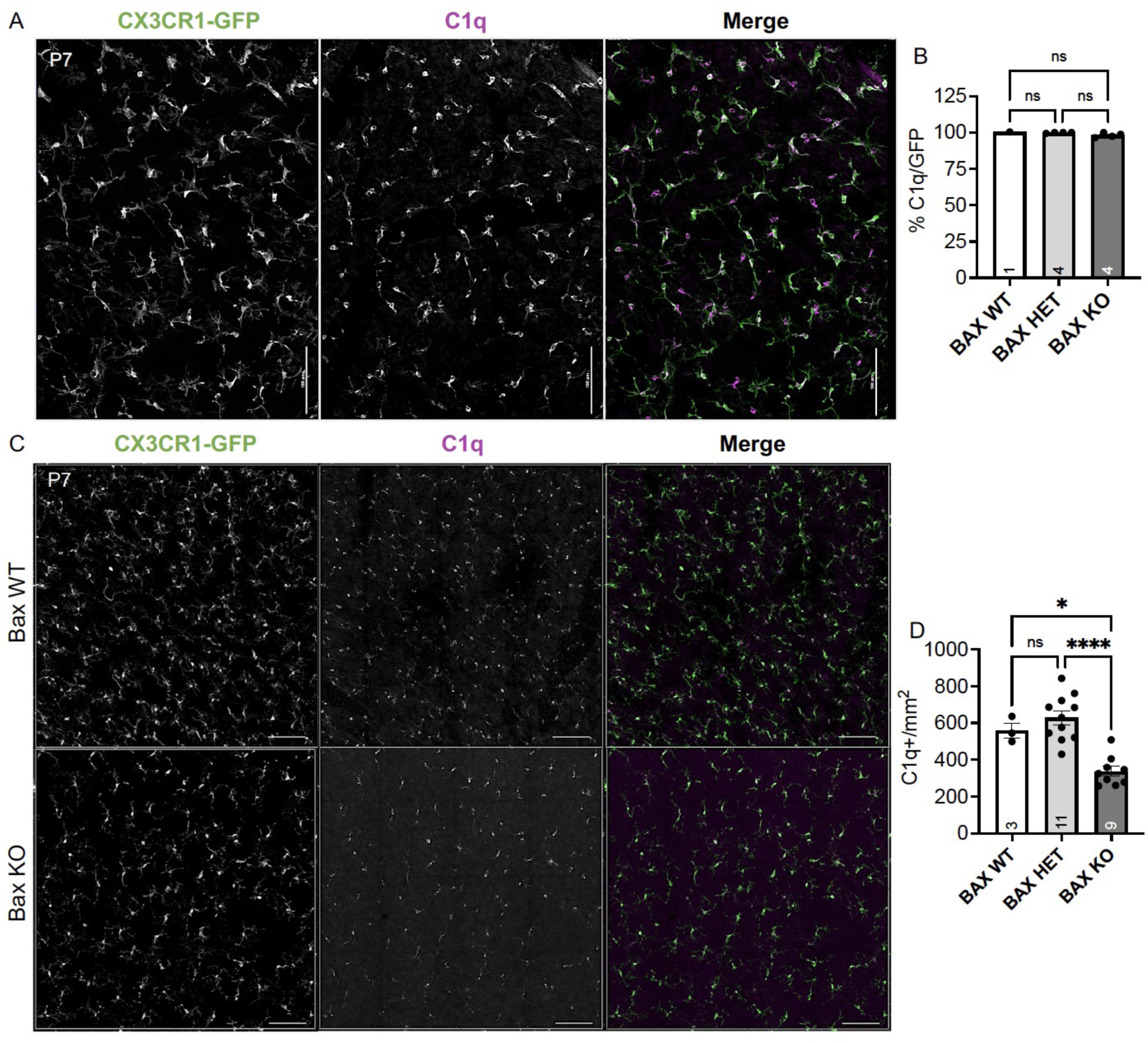
Bax KO retinas have reduced microglial density Related to Figure 3. (A) Max projected confocal image of a representative P7 whole mount retinas. CX3CR1-GFP/+ (green); C1q (magenta). Scale bar 100 µm. (B) Percent colocalization of C1q^+^ cells to CX3CR1-GFP^+^. (n=1 WT, n=4 HET, n=4 KO ; ±SEM) Kruskal-Wallis test statistic=4.818, p=0.0667. (C) Max projected confocal image of whole mount immunostained retinas of BAX WT and BAX KO. CX3CR1-gfp (green); C1q (magenta). Scale bar 50 µm. (D) Quantification of C1q^+^ microglia in a 0.526mm^2^ area of the dorsal, mid-peripheral, vascularized region of the retina. (n=3 WT, n=11 Het, n=9 KO; ±SEM) One-way ANOVA F(2,20)=19.90 p<0.0001 and Tukey’s multiple comparisons test *p=0.0112, ****p<0.0001.

**Figure S7.**
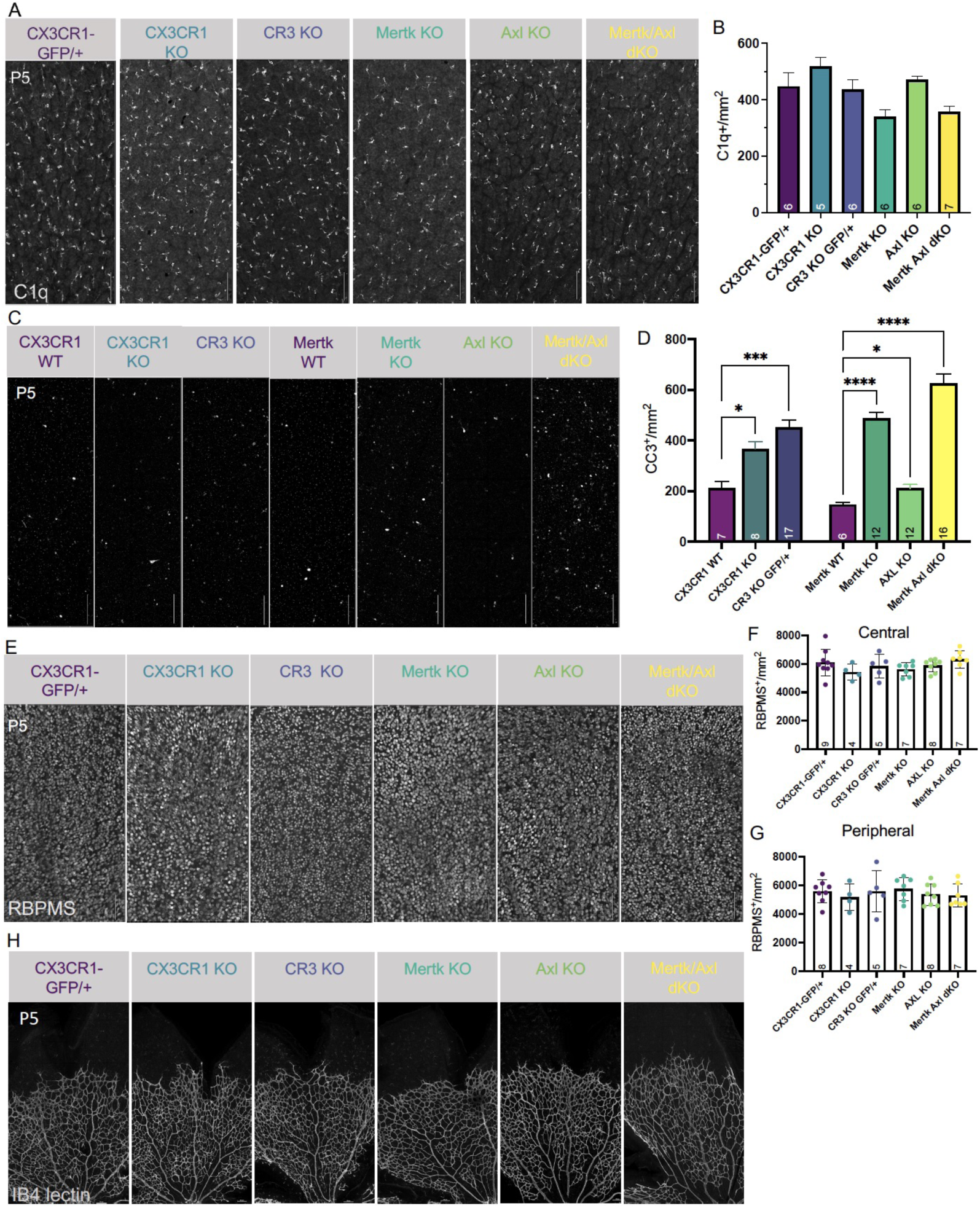
Loss of candidate receptors does not affect microglial density, RGC density, or BV development Related to Figure 5. (A) Max projected confocal images of whole mount immunostained retinas of all genotypes at P5 of dorsal leaf, mid-peripheral region. C1q (mono). Scale bar 100µm. (B) Microglia density (C1q^+^/mm^2^) for each genotype in dorsal leaf, mid-peripheral, vascularized region. (n=6 CX3CR1-GFP/+, n=5 CX3CR1 KO, n=6 CR3 KO CX3CR1-GFP/+, n=6 Mertk KO, n=6 Axl KO, n=7 Mertk Axl dKO; ±SEM). Kruskal-Wallis test statistic = 15.91 **p=0.0071 and Dunn’s multiple comparisons tests CX3CR1 KO vs Mertk KO *p=0.0489, CX3CR1 KO vs Mertk Axl dKO *p=0.0432, rest of the comparisons, ns. (C) Max projected confocal images of whole mount immunostained retinas of all genotypes at P5 of dorsal leaf, mid-peripheral region. CC3 (mono). Scale bar 100µm. (D) Total apoptotic body density (CC3^+^/mm^2^) for each genotype in entire dorsal leaf (Figure S9A). (n=7 CX3CR1 WT, n=8 CX3CR1 KO, n=17 CR3 KO CX3CR1-GFP/+, n=6 Mertk WT, n=12 Mertk KO, n=12 Axl KO, n=16 Mertk Axl dKO; ±SEM). Welch’s ANOVA W(6, 27.82)=54.71 ****p<0.0001 and Dunnett’s T3 multiple comparisons test. CX3CR1 WT vs CR3 KO ***p=0.0003, CX3CR1 WT vs CX3CR1 KO *p=0.0328, CX3CR1 WT vs Mertk WT ns, CX3CR1 WT vs Mertk KO ****p<0.0001, CX3CR1 WT vs Axl KO ns, CX3CR1 WT vs Mertk Axl dKO ****p<0.0001, CR3 KO vs CX3CR1 KO ns, CR3 KO vs Mertk WT ****p<0.0001, CR3 KO vs Mertk KO ns, CR3 KO vs Axl KO ****p<0.0001, CR3 KO vs Mertk Axl dKO *p=0.0257, CX3CR1 KO vs Mertk WT ***p=0.0007, CX3CR1 KO vs Mertk KO ns, CX3CR1 KO vs Axl KO *p=0.0089, CX3CR1 KO vs Mertk Axl dKO ***p=0.0004, Mertk WT vs Mertk KO ****p<0.0001, Mertk WT vs Axl KO *p=0.0467, Mertk WT vs Mertk Axl dKO ****p<0.0001, Mertk KO vs Axl KO ****p<0.0001, Mertk KO vs Mertk Axl dKO ns, Axl KO vs Mertk Axl dKO ****p<0.0001. Not all comparisons shown on graph. (E) Max projected confocal images of whole mount immunostained retinas of all genotypes at P5 of dorsal leaf, mid-peripheral region. RBPMS (mono). Scale bar 100µm. (F and G) RBPMS density for each genotype in dorsal leaf, central (F) and peripheral (G). (n=9,8 CX3CR1-GFP/+, n=4 CX3CR1 KO, n=5 CR3 KO CX3CR1-GFP/+, n=7 Mertk KO, n=8 Axl KO, n=7 Mertk Axl dKO; ±SEM). (F) Central: One-way ANOVA F(6,36)=1.251, p=0.3040 and (G) Peripheral: One-way ANOVA F(6,35)=0.4985, p=0.8051. (H) Max projected confocal images of whole mount immunostained retinas of all genotypes at P5 of dorsal leaf. IB4-lectin (mono). Scale bar 100µm.

**Figure S8.**
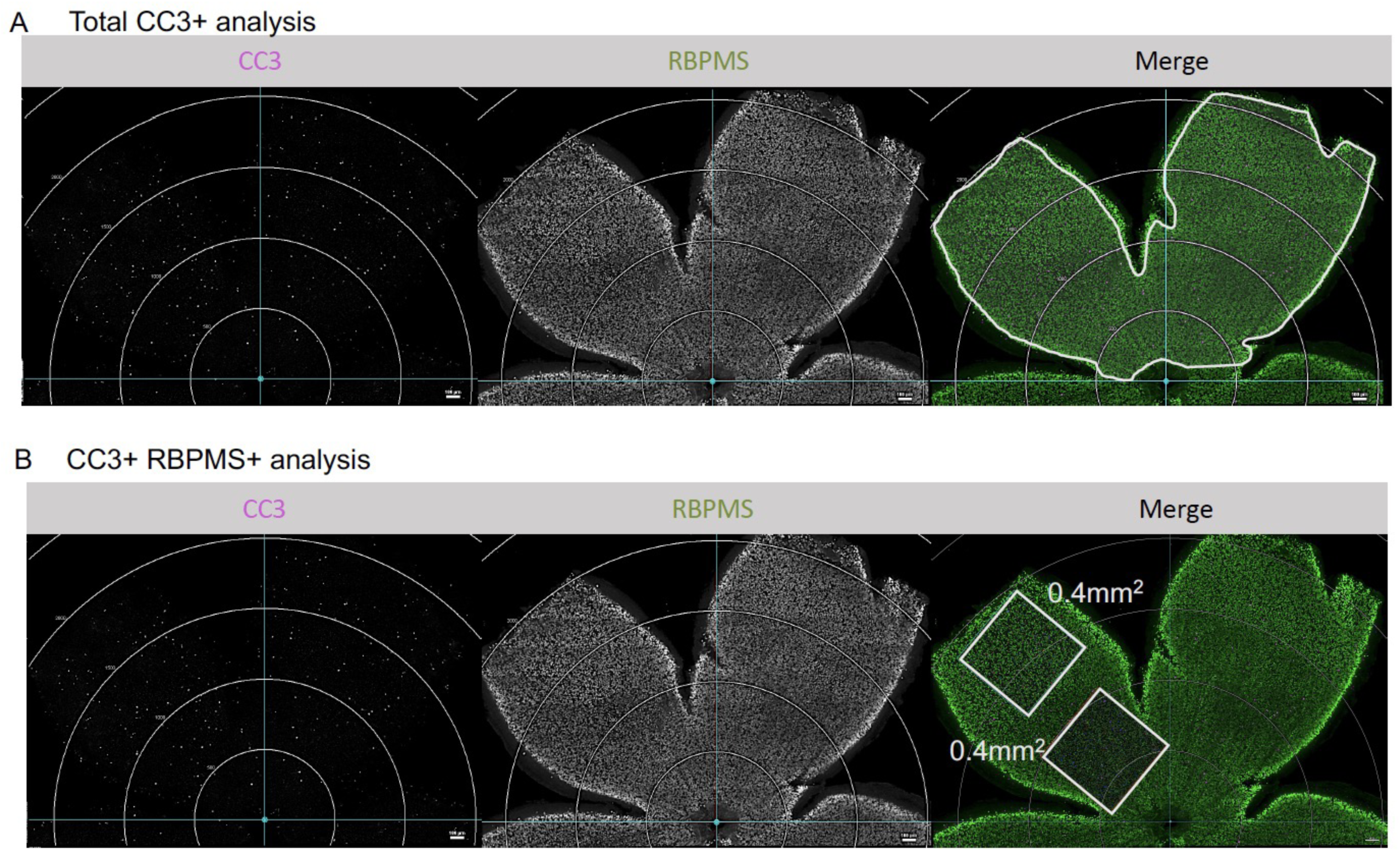
**Analysis of CC3+ and CC3+ RBPMS density Related to Figure 5 and S7**. (A) Max projected confocal images of whole mount immunostained retinas. Apoptotic cells, CC3 (magenta); RGCs, RBPMS (green). Circular grid in increments of 500µm. White line in merge outlines area analyzed for total CC3 analysis. Scale bar 100µm. (B) Confocal of whole mount immunostained retinas. Apoptotic cells, CC3 (magenta); RGCs, RBPMS (green). Circular grid in increments of 500µm. One central and one peripheral ROI of 0.400 mm^2^ was analyzed in the dorsal leaf and averaged. Scale bar 100µm.

**Figure S9.**
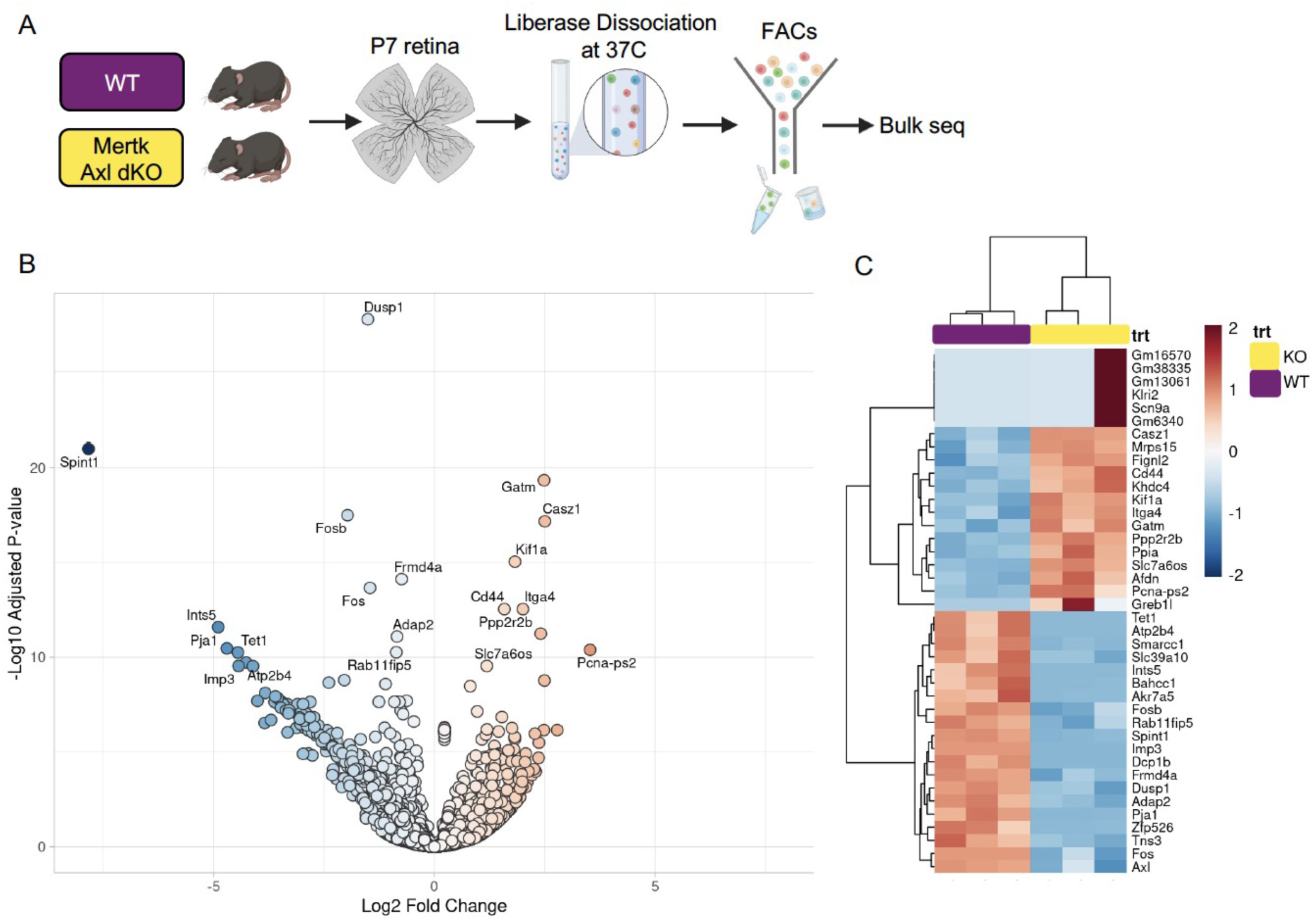
Mertk Axl dKOs do not have substantially altered lysosomal, lipid metabolism, or remodeling gene expression Related to Figure 7. (A) Workflow for P7 retinal collection, dissociation, sorting, and sequencing of individual microglia from 2 groups: WT, Mertk Axl dKO (n=3 each). (B) Volcano plot of differential expression analysis on WT compared to Mertk Axl dKO microglia. 404 upregulated and 1013 downregulated genes in dKO vs WT based on padj <0.05. (B) Heatmap of bulk sequencing on sorted microglia from WT and Mertk Axl dKO retinas top 20 genes upregulated and downregulated in dKO vs WT. Color represents z-score of normalized gene expression across all 6 samples.

## Notes

### Competing Interest Statement

The authors have declared no competing interest.

### Summary of Updates

Supplemental data tables are provided. Correction to the numbering of the supplemental figures.

